# Cryo-EM structures of photosystem I with alternative quinones reveals new insight into cofactor selectivity

**DOI:** 10.64898/2026.03.27.714801

**Authors:** Christian M. Brininger, Jimin Wang, Vasily Kurashov, Brandon P. Russell, Nikki Cecil M. Magdaong, David F. Iwig, Art van der Est, John H. Golbeck, David J. Vinyard, K. V. Lakshmi, Christopher J. Gisriel

**Author notes:** Current address: Department of Chemistry and Chemical Biology and The Baruch ‘60 Center for Biochemical Solar Energy Research, Rensselaer Polytechnic Institute, Troy, NY 12180.

## Abstract

Quinones are an integral component of electron transfer processes in photosynthetic and mitochondrial respiratory proteins. One such photosynthetic protein, Photosystem I, is an essential photooxidoreductase found in all oxygenic phototrophs. To better understand quinone chemistry and to form a basis for protein engineering, the *menB* gene in the model cyanobacterium *Synechocystis sp.* PCC 6803 was interrupted, blocking the biosynthesis of phylloquinone and causing it to be replaced by exchangeable plastoquinone-9 in the A_1A_ and A_1B_ quinone-binding sites of Photosystem I. This genetic variant has been instrumental in bioenergy research, enabling incorporation of a range of substituted and isotopically labeled quinones. Despite numerous valuable studies, the interpretation of biophysical data has been limited by a lack of structural data. To address this, we present the high-resolution cryo-EM structures of Photosystem I from the Δ*menB* variant containing (a) exchangeable plastoquinone-9 and (b) exogenously added 2-ethyl-1,4-napthoquinone at 1.90- and 2.05-Å resolution, respectively. Unexpectedly, the quinones in the A_1A_ and A_1B_ sites of Photosystem I, previously believed to have similar binding affinities, are found to be asymmetric in their ability to bind and exchange plastoquinone-9. This work reveals new and important insight into the molecular basis for Photosystem I activity in the Δ*menB* variant, the power of metabolic plasticity to maintain protein stability, and the requirement for protein instability to facilitate ligand exchange.

## Introduction

Photosynthesis is one of the most important biological reactions on Earth, being the primary source of energy for nearly all ecosystems^1^. In this solar powered process, energy from light is converted into chemical energy by transmembrane protein complexes known as reaction centers (RC)^2^. In the simplest view, all RCs contain two core homo- or hetero-dimeric membrane-spanning protein subunits that bind cofactors involved in electron transfer, always containing chlorophylls and/or bacteriochlorophylls and carotenoids^3^. By coupling light absorption to electron transfer through organic and metal-containing cofactors, reducing equivalents are generated to drive downstream metabolic processes.

There are two types of RCs: type I, in which the terminal electron acceptors are [4Fe-4S] clusters, and type II, in which quinone molecules are the terminal acceptors^4^. Type II RCs contain two bound quinones, one of which acts as a single-electron acceptor, while the other is a two-electron two-proton carrier that diffuses into the membrane as labile quinol following double reduction and protonation through proton-coupled electron transfer^5^. In contrast, type I RCs from plants, cyanobacteria, and green algae are known to bind two high-affinity quinones that act as single electron transfer intermediates, whereas type I RCs from anoxygenic organisms do not have tightly bound quinones participating in the electron transfer chain (ETC)^6,7^.

Oxygenic photosynthetic organisms contain both a type I RC, photosystem (PS) I ^8,9^, and a type II RC, PS II^5,10^, that have been extensively studied. The highly efficient light-driven electron transfer reactions of these photosystems are excellent blueprints for developing biohybrid and engineered systems for photocatalytic production of dihydrogen^11,12^ and carbon-based fuels^13–15^. Importantly, RCs more broadly are model systems for understanding energy and electron transfer reactions that are fundamental to the chemistry of life processes^16,17^. For example, it is known that the use of both type I and type II RCs in oxygenic photosynthesis provides organisms with a major competitive advantage by generating a highly oxidizing potential in PS II, so that water can be used as an electron source, while at the same time producing a very negative reduction potential in PS I, which is needed for the generation of NADPH^18^.

Cyanobacterial PS I is an approx. 1 MDa protein complex consisting of 10 - 12 protein subunits and >100 ligands^19^. The ETC of PS I (**Fig. 1*A***) begins with a pair of excitonically coupled chlorophyll (Chl) *a*/*a*′ molecules P_A_ and P_B_ that form the donor P_700_. Subsequently, the ETC bifurcates into two branches, with each branch containing two Chls, A_–1(A or B)_ and A_0(A or B)_, and a quinone in the A_1(A or B)_ sites^20–22^. The two branches converge at the [4Fe-4S] cluster F_X_, which is followed by two terminal [4Fe-4S] clusters, F_A_ and F_B_^23^. An intriguing aspect of PS I is the presence of high affinity quinones, typically phylloquinone (PhQ), in the A_1(A or B)_ sites. The redox chemistry of oxygenic photosynthesis shows that PS I evolved to produce a highly negative reduction potential and retained these quinones even though they are not typically strong reductants. Interestingly, anoxygenic type I RCs, considered representative of primitive evolutionary precursors of PS I, are not as strong reductants and lack bound quinones^6,7,24^.

**Fig. 1.**
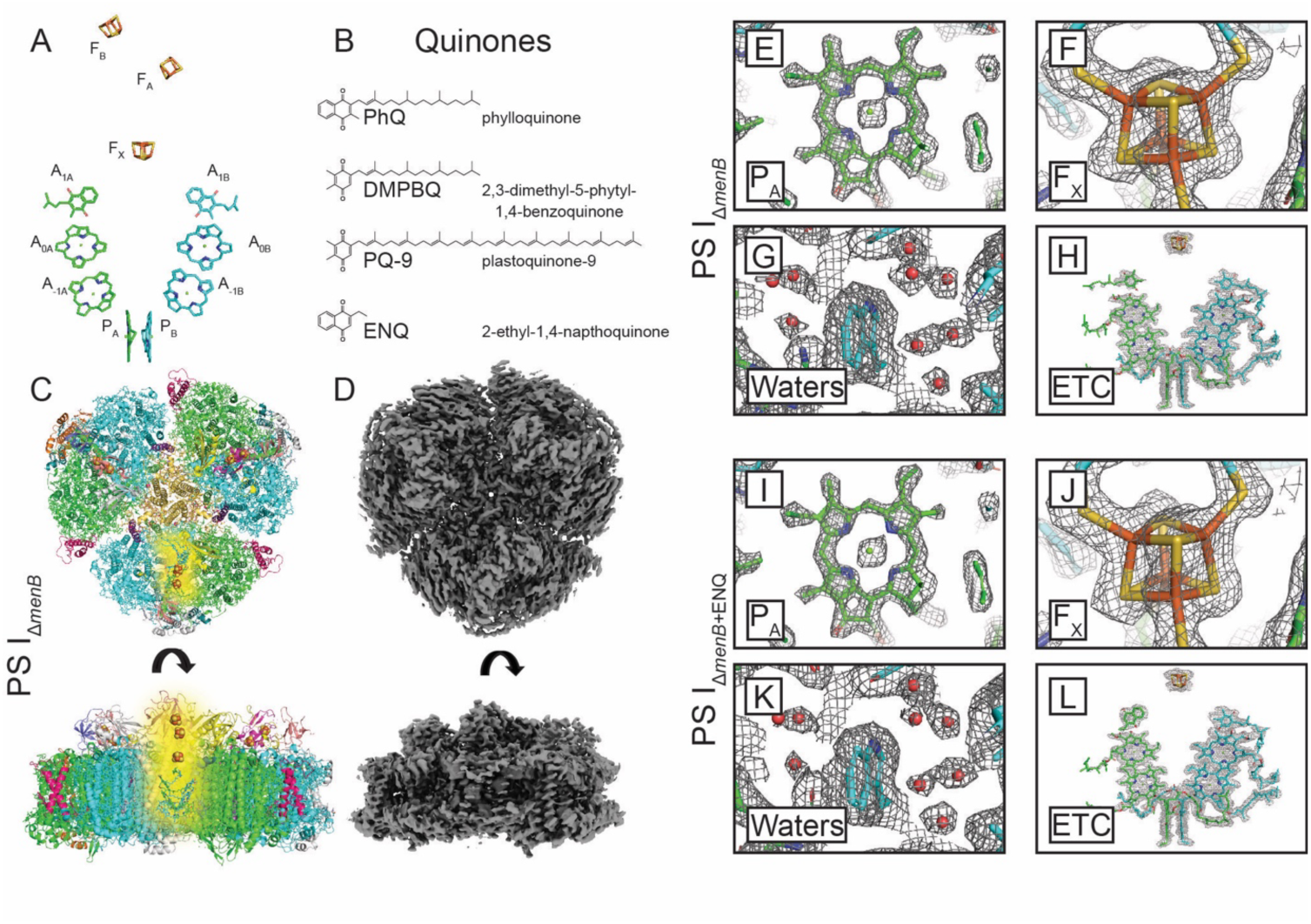
PS I electron transfer chain, relevant quinones, and cryo-EM images. (*A*) Native PS I ETC cofactors. Tails and ring substituents are hidden for clarity. (*B*) Quinones relevant to the work presented herein. (*C*) Model of the PS IΔ*menB* structure with ETC cofactors highlighted in one monomeric unit. (*D*) Unsharpened map of the PS IΔ*menB* structure. (*E-H*) Example regions of the PS IΔ*menB* structure. (*I-L*) Example regions of the PS IΔ*menB*+ENQ structure.

To identify the genes involved in the biosynthesis of PhQ and generate a system for engineering PS I to perform alternative chemistries, a series of variants were previously constructed in which genes in the putative biosynthetic pathway were disrupted in *Synechocystis* sp. PCC 6803 (hereafter *Synechocystis*)^25–27^. In two of these variants, Δ*menA* and Δ*menB*, the synthesis of PhQ was interrupted but photosynthetic growth was inhibited only under high light conditions. Analysis of the ETC cofactors in PS I from these variants revealed the incorporation of plastoquinone-9 (PQ-9) and absence of the native PhQ (**Fig. 1*B***)^28^, the former of which is the quinone type involved in PS II electron transfer^29^. Moreover, the loosely bound PQ-9 in PS I from the Δ*menB* variant was replaced *in vitro* upon incubation with exogenous naphthoquinones. This made it possible to probe the kinetics and thermodynamics of electron transfer through quinone replacement in the Δ*menA* and Δ*menB* variants.

Although there exists a vast body of biophysical studies on PS I from the Δ*menB* variant, there is a lack of high-resolution structural data at this time. Since the Δ*menB* variant was first constructed more than 20 years ago^25^, several related strains of the *ΔmenB* variant have emerged. In this work, we will refer to three *Synechocystis* variants: *ΔmenB(2002)*^25^, which is the original strain maintained as a frozen stock that contains PS I with displaceable PQ-9; *ΔmenB(2023)*^30,31^, the original strain intermittently maintained as a living culture that contains PS I with 2,3-dimethyl-5-phytyl-1,4-napthoquinone (DMPBQ) as a result of evolution under laboratory conditions; and *ΔmenB(2024)*^31^, a newly generated strain that is the topic of this study. Recently, we reported the structure of PS I from the *ΔmenB(2023)* variant that contained high-affinity DMPBQ molecules in both A_1_ sites^30^. Because the phenotype and quinone exchangeability of the *ΔmenB(2023)* variant resembled the wild-type strain over time, it was proposed that the presence of DMPBQ was a consequence of “laboratory evolution”: the cells had evolved to incorporate the non-exchangeable, and hence more stable DMPBQ that possesses a nineteen carbon atom (C-19) tail, which is identical to the native PhQ rather than the exchangeable PQ-9 that possesses a 45 carbon atom (C-45) tail. This was confirmed by whole genome analysis of the *ΔmenB(2023)* variant revealing a mutation in the tocopherol cyclase gene (*slr1737*) in the tocopherol biosynthesis pathway^31^. The *ΔmenB(2023)* strain sacrificed tocopherol synthesis, the absence of which is known to have no effect on growth under laboratory conditions, resulting in incorporation of DMPBQ in the A_1A_ and A_1B_ sites. Hereafter, we refer to PS I from the *ΔmenB(2023)* laboratory-evolved strain as PS I_evolΔ*menB*_. Although that study provided insight into laboratory evolution and quinone stabilization, it was not possible to use that strain to obtain structural data on PS I containing exogenously added quinones in the A_1A_ and A_1B_ sites as DMPBQ has a high binding affinity and is, therefore, non-exchangeable^31^.

Here, we report the structural characterization of PS I from a newly generated Δ*menB(2024)* deletion strain (hereafter referred to as PS I_Δ*menB*_). This strain is a recapitulation of the original *ΔmenB(2002)* strain in which growth occurs under low light and quinone exchange has been recovered^31^. Cryo-electron microscopy (cryo-EM) was used to determine single-particle reconstructions of PS I from PS I_Δ*menB*_ without and with the incubation of exogenously added 2-ethyl-1,4-naphthoquinone (ENQ) (the latter of which is hereafter referred to as PS I_Δ*menB+*ENQ_). The structures of these PS I complexes show that the A_1A_ and A_1B_ sites exhibit different affinities and varied degrees of incorporation of PQ-9 and DMPBQ in PS I*_τιmenB_*. Moreover, PQ-9 in these sites can be displaced by exogenously added ENQ due to the displacement of nearby ligands and peripheral subunits. These data provide a structural basis for interpretation of previous biophysical analyses and reveal novel observations about quinone stability and its relevance to fundamental aspects of PS I structure.

## Results

### Sample preparation, characterization, and cryo-electron microscopy

The Δ*menB(2024)* variant was generated and propagated as described in Russell et al^31^. In brief, the *menB* gene was interrupted using insertional inactivation and verified using whole genome sequencing. PS I_Δ*menB*_ was purified as summarized in **Materials and Methods**. The absorption spectrum of isolated PS I_Δ*menB*_ was consistent with typical cyanobacterial WT PS I complexes, with peak maxima at 680, 595, and ∼440 - 385 nm corresponding to the Q_y_, Q_x_, and Soret transitions of Chl *a*, respectively (*SI Appendix*, **Fig. S1**). Liquid chromatography/mass spectrometry (LC/MS) employing a 1:1 mixture of methanol/acetone extracts of PS I_Δ*menB*_ and the WT were used to characterize the respective quinone species. The LC/MS ion chromatogram of WT PS I (*SI Appendix*, **Fig. S2**) displayed a major species that elutes at ∼8.6 min with an *m/z* of 451.3571, which is consistent with PhQ. This species is not present in PS I_Δ*menB*_, which contains DMPBQ and PQ-9 instead, that elute at ∼7.75 min and ∼9.25 min with an *m/z* of 415.3571 and 749.6231, respectively. Similarly, pigment analysis using HPLC detected plastoquinone (corresponding to PQ-9) and naphthoquinone (corresponding to ENQ) in the PS I_Δ*menB*_ sample (*SI Appendix*, **Fig. S3**).

Quinone exchange with exogenously added ENQ to generate PS I_Δ*menB*+ENQ_ is described in **Materials and Methods**. Relative to the PS I_Δ*menB*_ sample, pigment analysis detected an additional peak corresponding to ENQ (*SI Appendix*, **Fig. S3**). For preliminary structural characterization, the samples were negatively stained and imaged using transmission electron microscopy, revealing monodisperse trimeric PS I particles (*SI Appendix*, **Fig. S4**). To generate higher resolution structural data, the samples were subjected to single particle cryo-EM. Initial screening confirmed monodispersity of trimeric PS I particles (*SI Appendix*, **Fig. S5**) and high-resolution datasets were collected. After single-particle analysis, the final cryo-EM maps exhibited global resolutions of 1.90 Å and 2.05 Å for PS I_Δ*menB*_ and PS I_Δ*menB*+ENQ_, respectively. Example micrographs, 2D classifications, workflows, and data statistics are shown in *SI Appendix*, **Fig. S6-S8**, and **Table S1**. The model and map for the PS I_Δ*menB*_ structure are shown in **Fig. 1*C*** and **1*D***, and example map regions from the PS I_Δ*menB*_ and PS I_Δ*menB*+ENQ_ structures are shown in **Fig. 1*E-L***.

### Quinone tail analysis

We analyzed the tail regions to obtain insight into the quinone species bound in the A_1A_ and A_1B_ sites of the PS I_Δ*menB*_ and PS I_Δ*menB*+ENQ_ maps and compared these to the recent cryo-EM data of PS I_evolΔ*menB*_ from the *ΔmenB(2023)* strain^30^ (**Fig. 2**). Relevant quinones in this analysis are shown in **Fig. 1*B***. PQ-9 contains a nine-isoprenoid unit C-45 tail and DMPBQ has a C-19 phytyl tail with a single isoprenoid group. For the PS I_Δ*menB*_ structure, the map signal for the quinone tail in the A_1A_ site was observed to approx. the 10^th^ carbon atom along the tail, after which the signal diminished in intensity and is less defined. Additionally, the map corresponding to other features from the surrounding amino acids and ligands in the vicinity of the quinone in the A_1A_ site were also poorly resolved in comparison to the rest of the map and to the PS I_evolΔ*menB*_ structure, suggesting inherent disorder. This includes the PsaF and PsaJ subunits, β-carotenes J13 and J12, Chl F1, and the tails of A1, A38, A39, A40 Chl molecules (*SI Appendix*, **Fig. S9**, nomenclature is based on Jordan et al. 2001^3^). Based on these observations, we propose that the A_1A_ site contains primarily PQ-9, and that its long flexible tail leads to disorder in the surrounding environment, making it difficult to resolve the tail in its entirety.

**Fig. 2.**
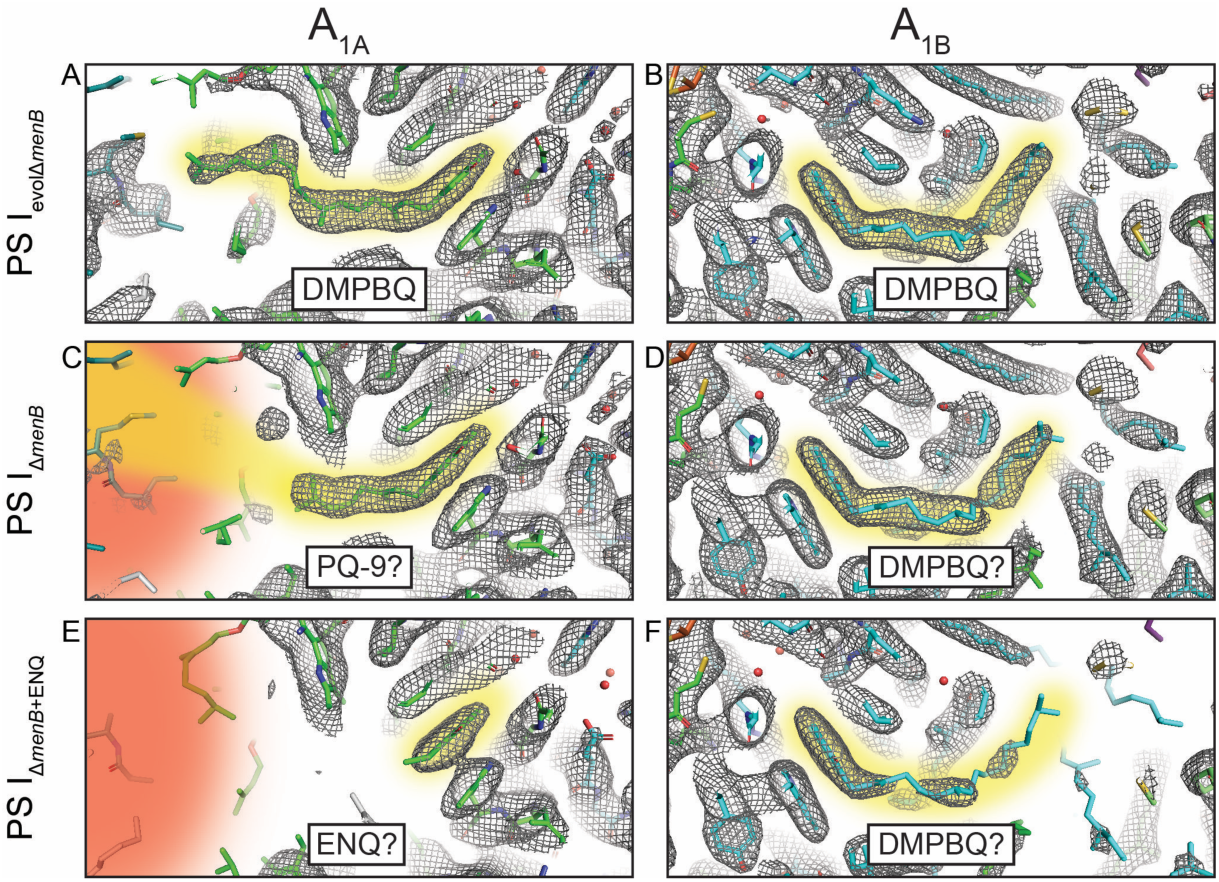
PS I A1A and A1B quinone tail densities. Structural models are shown within the unsharpened cryo-EM maps, with the modeled quinones highlighted in yellow. (*A*, *B*) Model and map of the A1A and A1B sites for PS IevolΔ*menB* (PDB: 9AU4), respectively. (*C*, *D*) Model and map of the A1A and A1B sites for PS IΔ*menB*, respectively. (*E*, *F*) Model and map of the A1A and A1B sites for PS IΔ*menB*+ENQ, respectively. The red shaded area depicts the disordered regions in the respective binding pockets.

The tail of the quinone in the A_1B_ site, on the other hand, more closely resembles the PS I_evolΔ*menB*_ structure which contains DMPBQ (C-19 tail). Although the tail is slightly less well resolved (*SI Appendix*, **Fig. S10**), it appears to be the same length (**Fig. 2**), implying the site is occupied primarily by DMPBQ. Supporting this hypothesis, the tail appears generally inaccessible to the solvent, being buried by nearby protein features including residues from PsaB, β-carotenes B17 and L19, Chls A31 and B39, and the galactolipid in site B2, all of which have well-defined density in the cryo-EM map, similar to that of PS I_evolΔ*menB*_.

Because quinone tail length appears to influence the protein stability, we visualized the PS I_evolΔ*menB*_, PS I_Δ*menB*_, and PS I_Δ*menB*+ENQ_ structures using Q-scores (**Fig. 3**), which quantitate the fit of structural coordinates within a cryo-EM map^32^. Note that the Q-score generally decreases from the center of the complex, where proteins are usually less flexible resulting in higher local resolution, to the periphery of the complex, where proteins are more flexible resulting in lower local resolution. On the B-side of the complex in the vicinity of the quinone tail in the A_1B_ site, where the structural features of PS I_Δ*menB*_ resemble those of the PS I_evolΔ*menB*_ structure (**Fig. 2**), the Q-scores are highly similar among the structures compared. Notably, the A_1B_ site is situated close to the center of the PS I trimer near the oligomerization interface where the quinone tail may be relatively constrained in its position and therefore may exhibit more stability. However, Q-scores in the vicinity of quinone tail in the the A_1A_ site are much lower in the PS I_Δ*menB*_ structure compared to that of PS I_evolΔ*menB*_, implying greater flexibility in the former. The A_1A_ site is located towards the periphery of the PS I complex, close to the PsaF and PsaJ subunits, and regions of the PsaA and PsaB core subunits (**Fig. 3**). Based on this analysis, we hypothesize that the A_1A_ site of the PS I_Δ*menB*_ structure is at least partially occupied by PQ-9 (C-45 tail), which leads to disorder in the surrounding protein environment. The A_1B_ site, however, seems likely to be primarily occupied by DMPBQ (C-19 tail), and exhibits more order. If this is the case, it suggests that the A_1A_ site would be much more amenable to quinone exchange than the A_1B_ site.

**Fig. 3.**
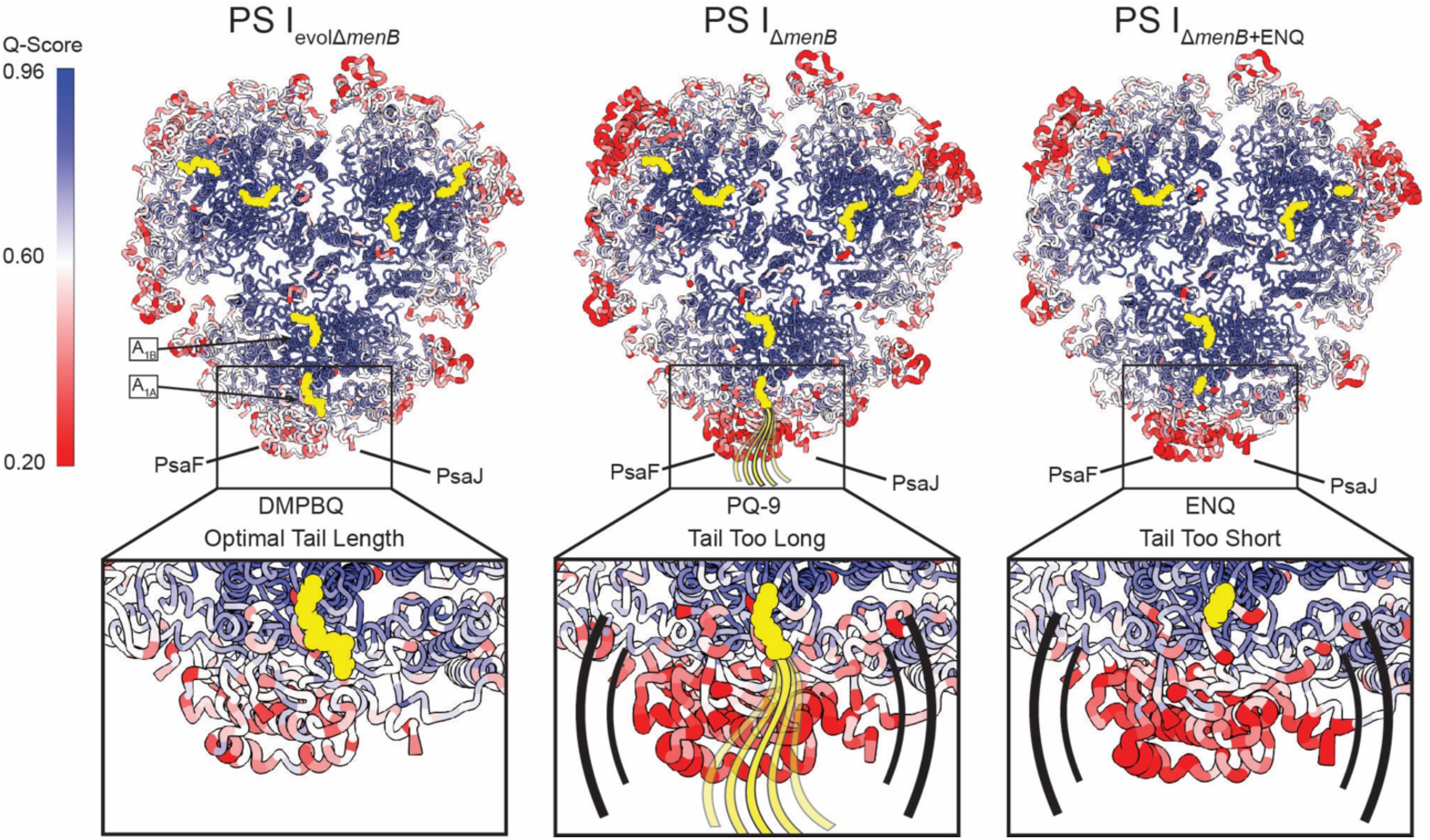
Structural instability comparison based on Q-score. The structures of PS IevolΔ*menB*, PS IΔ*menB*, and PS IΔ*menB*+ENQ are colored by their Q-score values. High Q-score corresponds to higher protein resolvability, and lower Q-score corresponds to lower protein resolvability, implying more or less stability, respectively. The quinone binding sites A1A and A1B are highlighted in yellow in the PS IevolΔ*menB* structure only.

For the PS I_Δ*menB*+ENQ_ structure, where PS I_Δ*menB*_ was incubated with exogenously added ENQ, the map for the quinone occupying the A_1A_ site displayed minimal density for the tail (**Fig. 2** and *SI Appendix*, **Fig. S11**), which is expected as the site is occupied by ENQ. This supports the hypothesis that the A_1A_ site in the PS I_Δ*menB*_ structure is primarily occupied by PQ-9, and upon incubation with ENQ, is exchanged due to the higher binding affinity of ENQ^33^. In the Q-score analysis, the region surrounding the A_1A_ site in the PS I_Δ*menB*+ENQ_ structure also exhibits lower Q-scores than the PS I_evolΔ*menB*_ structure, similar to the PS I_Δ*menB*_ (**Fig. 2**). We propose that the lack of a tail in ENQ, or the presence of a long C-45 tail in PQ-9, results in disorder in the surrounding protein environment. These observations are unlike the A_1B_ site in the PS I_Δ*menB*+ENQ_ structure, where the quinone tail region appears somewhat more ambiguous, but is similar to both the corresponding sites from the PS I_evolΔ*menB*_ and PS I_Δ*menB*_ structures (*SI Appendix*, **Fig. S10** and **S11**). This supports the hypothesis that the high affinity quinone DMPBQ is present at substantial occupancy in the A_1B_ site of the PS I_Δ*menB*+ENQ_ structure, which is non-exchangeable^31^.

### Quinone headgroup analysis

We next analyzed the quinone headgroup regions in the A_1A_ and A_1B_ site of the PS I_Δ*menB*_ structure (**Fig. 4**). The headgroup of both PQ-9 and DMPBQ is benzoquinone, which exhibits a single conjugated six-membered ring, whereas PhQ and ENQ are naphthoquinones, with two fused conjugated six-membered rings (**Fig. 1*B***). The headgroup of the quinones in both sites of the PS I_Δ*menB*_ structure are benzoquinones. This is expected, as there should be only two quinone assignments possible: DMPBQ or PQ-9. The situation is more complex for the PS I_Δ*menB*+ENQ_ structure. In the A_1A_ site, the headgroup density clearly fits a naphthoquinone, which is consistent with the observation that the site contains ENQ at high occupancy based on the tail analysis (*SI Appendix*, **Fig. S11**). The ethyl moiety is observed primarily in one position at high threshold, however, a second or alternate orientation is apparent at lower threshold (*SI Appendix*, **Fig. S12**). The cryo-EM maps are ensemble data derived from hundreds of thousands of individual particle projections, therefore, this observation suggests that although the A_1A_ site contains ENQ at high occupancy, it does so in two possible orientations.

**Fig. 4.**
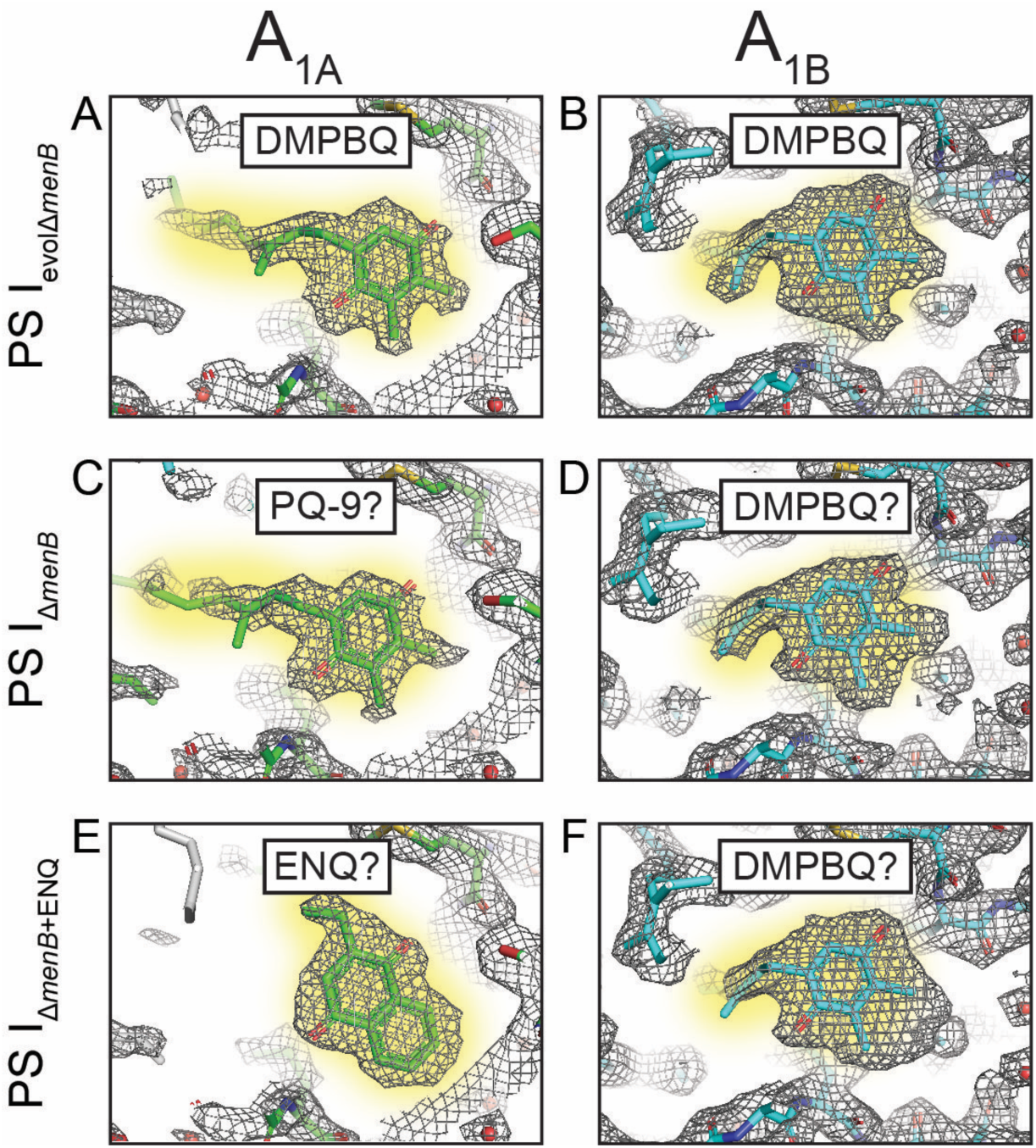
A1A and A1B quinone headgroup densities in PS. **I.** Structural models are shown within the sharpened maps with the quinones highlighted in yellow. (*A*, *B*) Model and map for A1A and A1B in PS IevolΔ*menB* (PDB: 9AU4). (*C*, *D*) Model and map for A1A and A1B quinones in PS IΔ*menB*. (*E*, *F*) Model and map for A1A and A1B quinones in PS IΔ*menB*+ENQ.

To deconvolute the occurrence of each orientation, we developed a procedure (*SI Appendix*, **Text S1** and **Fig. S13**) to estimate the occupancies by sampling the map intensity along selected carbon atoms following the geometry of the naphthoquinone headgroup (**Fig. 5A**). The analysis allowed us to estimate that ENQ in a “flipped” orientation (relative to the orientation of the tail of the PhQ in WT PS I) represents a majority population (∼75%) and ENQ in an “unflipped” orientation (the same orientation as the PhQ tail in WT PS I) represents a minority population (∼25%) (**Fig. 5B**).

**Fig. 5.**
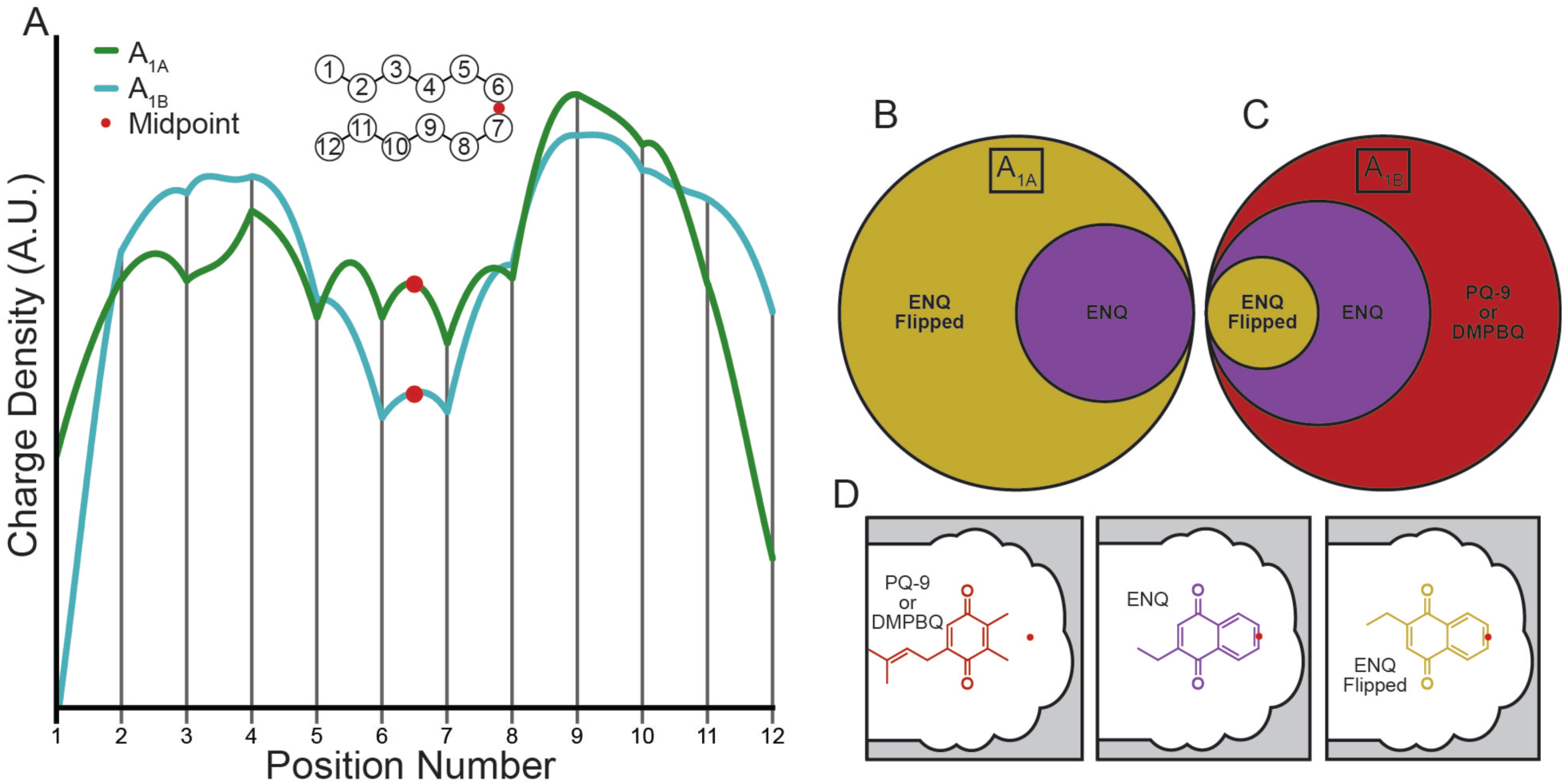
Charge density analysis of quinone site occupancy in PS IΔ*menB*+ENQ. (*A*) Charge density traces for A1A and A1B sites. (*B,C*) Representation of approximate occupancies in the A1A and A1B sites respectively. (D) Legend showing the orientations/identities of the modeled species.

It is noteworthy that two previous transient electron paramagnetic resonance (TR-EPR) spectroscopy studies in literature also indicated that ENQ binds primarily in a flipped orientation to the A_1A_ site of PS I^33,34^. In the first study^27^, the PhQ was solvent-extracted from WT PS I and, replaced by ENQ. The resulting TR-EPR spectrum, which arises exclusively from the P ^+^A ^−^ _700 1A_ radical pair, displayed strong electron-nuclear hyperfine couplings to the methylene protons of the ethyl side chain, indicating higher electron spin density on the adjacent ring carbon atom. The X-ray structure of native WT PS I^3^ and HYSCORE spectroscopy studies^35^ show that phylloquinone is involved in a single H-bond and that the H-bonded carbonyl group is ortho to the phytyl tail and meta to the methyl group. The single H-bond distorts the electron spin density distribution in the semiquinone radical, resulting in higher spin density on the carbon adjacent to the methyl group and lower spin density adjacent to the phytyl tail. This leads to strong hyperfine coupling to the methyl protons and weak coupling to the methylene protons of the phytyl tail^34^. Thus, the strong hyperfine coupling to the methylene protons observed for ENQ in the A_1_ site led to the conclusion that the ethyl moiety was in the position normally occupied by the methyl group of PhQ in WT and not in the position of the phytyl tail, i.e. the quinone was in a flipped orientation. A subsequent TR-EPR study performed on PS I_Δ*menB*+ENQ_ from the Δ*menB(2002)* strain^33^ showed that the spectrum for the latter was similar to that observed in the previous study that incorporated ENQ into solvent-treated PS I, supporting the assignment of the flipped orientation in the A_1A_ site. Indeed, systematic studies of a series of quinones bound to the A_1A_ site showed a very strong tendency to incorporate quinones with a substituent meta to the single H-bonded carbonyl group and if present, a methyl substituent would preferentially occupy that position. We performed TR-EPR on the PS I_Δ*menB*_ and PS I_Δ*menB*+ENQ_ samples from the Δ*menB(2024)* strain in this study (**Fig. 6**), comparing it to the previous TR-EPR data in literature^33^. The spectra obtained from the Δ*menB(2002)* and Δ*menB(2024)* strains incubated with ENQ are virtually identical, demonstrating the high degree of reproducibility of the spectrum of PS I_Δ*menB*+ENQ_. Together with the cryo-EM data, this elegantly demonstrates that the unique TR-EPR spectrum of PS I with bound ENQ, first observed more than two decades ago, arises due to the majority population of ENQ in a flipped orientation in the A_1A_ site.

**Fig. 6.**
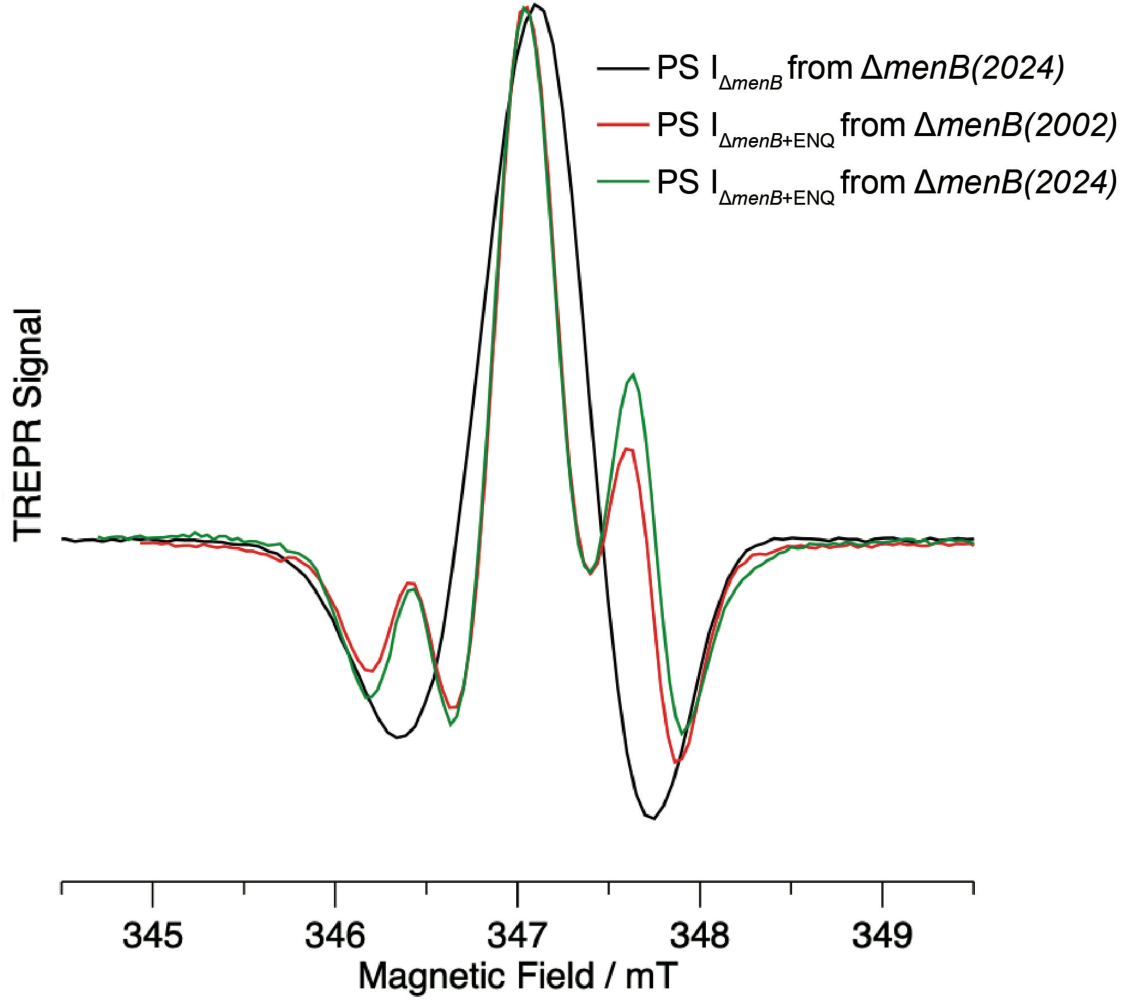
TR-EPR spectroscopy of PS I samples. The TR-EPR spectra of PS IΔ*menB* from the Δ*menB(2024)* strain (black) and PS IΔ*menB*+ENQ from the Δ*menB(2024)* (green) and Δ*menB(2002)* (red) strains^33^. Spectra are normalized to the same maximum amplitude. A linear baseline correction was applied to the PS IΔ*menB* spectrum from the Δ*menB*(2024) strain to remove the broad contribution arising from the spectrum of the triplet state of P700 formed by recombination from A0^−^.

The A_1B_ site of PS I_Δ*menB*+ENQ_ is more complex than the A_1A_ site. The headgroup density in the cryo-EM map appears to fit a benzoquinone with a single conjugated ring at higher threshold, but an additional signal for the larger naphthoquinone headgroup with two fused conjugated rings becomes visible at lower threshold (**Fig. 4** and *SI Appendix*, **Fig. S12**). We interpret this as the A_1B_ site containing a mixed occupancy of ENQ and PQ-9 or DMPBQ. We performed an analysis of the cryo-EM map which revealed three populations: PQ-9 or DMPBQ (which have the same benzoquinone headgroup) is the majority population at ∼66%, ENQ in the unflipped orientation is a smaller population at ∼25%, and ENQ in the flipped orientation is a very small minority at ∼10% (**Fig. 5B**). These results expand upon the quinone tail analysis, showing that while the majority of the A_1B_ sites in PS I_Δ*menB*_ are likely occupied by DMPBQ, there is a minority fraction of sites containing PQ-9 that can be exchanged with ENQ. This explains why the A_1B_ tail of PS I_Δ*menB*_ is slightly less well resolved than the corresponding density in the PS I_evolΔ*menB*_ structure (*SI Appendix*, **Fig. S10**).

## Discussion

It has been shown that at low temperature, the TR-EPR signals of PS I_Δ*menB*_ incubated with naphthoquinones arise solely from P_700_^+^A_1A_^− 36^. Thus, such data does not provide direct information about the occupancy of the A_1B_ site. As a result, in the absence of any indication to the contrary, it was assumed that the quinones in both A_1A_ and A_1B_ are exchangeable, given the similarity of the binding sites^25^. Perhaps the most striking insight provided by the present study is that whereas the A_1A_ site of PS I_Δ*menB*_ appears fully exchangeable, the A_1B_ site exhibits different fractional occupancies of exchangeable and non-exchangeable quinones. Before drawing further conclusions, it is important to note that the cryo-EM occupancy analysis presented here may be limited by two important factors. First, it does not account for empty sites, i.e. it is limited to analysis of the fractions of occupied sites. The TR-EPR spectrum of PS I_Δ*menB*_ isolated from the Δ*menB(2024)* strain, however, exhibits a P_700_ triplet state signal (*SI Appendix*, **Fig. S14**) which arises only when A_1_ sites are not occupied. Quantifying the size of the fraction of empty A_1_ sites in the PS I_Δ*menB*_ from the TR-EPR data is not straightforward, but following incubation with ENQ, the triplet P_700_ signal is absent (*SI Appendix*, **Fig. S14**). This implies that all empty A_1_ sites have been filled with ENQ. Second, the cryo-EM occupancy analysis assumes that the headgroups of the different quinones are approximately aligned. In support of this assumption, pulsed EPR studies suggest that the P_700_ - A_1A_ distance is the same (within 0.3 Å) comparing WT PS I to (a) PS I_Δ*menB*_ isolated from the Δ*menB(2002)* strain and (b) PhQ-depleted PS I (through solvent extraction) after incubation with four different quinones.^37^ Furthermore, the polarization patterns of the TR-EPR spectra shows that the carbonyl groups of the quinones are all aligned approximately parallel to the vector connecting P_700_^+^ and A_1_^− 33^. Despite this, minor differences in the positions of the headgroups comprising the ensemble cryo-EM map are possible and could influence the determination of occupancies. Similarly, we avoid a thorough analysis of the headgroup interactions (e.g. H-bonding), as the error involved in modeling the atomic positions is challenging to account for on the scale of a few tenths of an Å, especially considering fractional occupancy. However, the hyperfine coupling patterns in the TR-EPR spectra suggest asymmetric H-bonding interactions. Detailed insights on the H-bonding interactions will be provided in the future through computational modeling based on the atomic coordinates provided by the fits to the cryo-EM maps and hyperfine correlation spectroscopy.

In the A_1B_ site of PS I_Δ*menB*_, the evidence for fractional occupancy of PQ-9 is based on the lower definition of the tail density relative to the PS I_evolΔ*menB*_ structure, yet after incubation with ENQ, we find that ∼35% of the A_1B_ sites contain ENQ (**Fig. 5**). Based on the structure of PS I, it is difficult to envision how PQ-9 might fit, or exchange from this site. This is because it would require rearrangement of the surrounding protein environment involving at least ∼30 N-terminal residues of PsaB, ∼5 C-terminal residues of PsaI, and numerous cofactors including Chl B38, β-carotene L19, and the galactolipid in site B2 (*SI Appendix*, **Fig. S15**). Yet, there are no substantial differences in the map quality in any of these regions in the PS I_Δ*menB*_ map compared to the PS I_evolΔ*menB*_ map. It is possible that ∼35% of the sites being occupied by PQ-9 and the associated protein flexibility is too low to detect differences in that region. However, given the spectroscopic evidence for charge recombination between P_700_^+^ and A_0_^−^, it seems most likely that the A_1B_ sites containing ENQ are unoccupied prior to incubation. Clearly, more work is needed to conclusively determine why a fraction of ENQ is observed in the PS I_Δ*menB*+ENQ_ A_1B_ site without corresponding structural features in the PS I_Δ*menB*_ map. Despite this uncertainty, we can conclude that the occupancy of the A_1A_ and A_1B_ sites in the Δ*menB* strain differ and the A_1A_ site is much more likely to be solely occupied by PQ-9, than the A_1B_ site.

The asymmetry in quinone occupancy in the A_1A_ and A_1B_ sites was a surprising result and raises intriguing questions about the assembly of the PS I complex. It may also provide more fundamental insight into the role of quinones in RCs. It has been previously shown that DMPBQ binds tightly the the A_1A_ site when it is present and it cannot be displaced even by native PhQ^31^. While the cryo-EM structures indicate that the A_1B_ site of PS I_Δ*menB*_ contains DMPBQ, it is puzzling how PS I complexes with DMPBQ bound only to the A_1B_ site can be formed. A_1B_ is close to the center of the PS I trimer, near the monomer-monomer interfaces. That location probably imposes a more compact environment. One could imagine that during PS I biogenesis, some monomers bind whatever small fraction of DMPBQ is available more readily to the A_1B_ site, stabilizing the complex so that trimerization is not disrupted. Those that contain PQ-9 in A_1B_, however, would be less stable, causing the fraction of successful trimerization events to decrease. To test this hypothesis, we performed blue native gel electrophoresis of solubilized thylakoid membranes (*SI Appendix*, **Fig. S16**). The results qualitatively show that the PS I monomer : trimer ratio is higher in *ΔmenB(2024)* compared to the WT, supporting the hypothesis that non-optimal tail length in the A_1B_ site decreases the likelihood of PS I trimerization. Such instability is less consequential on the A-side near the periphery of PS I; PQ-9 can bind at high occupancy without disruption of trimerization, and thus A_1A_ is more exchangeable. This general scheme is also observed in native PS II^38^, where the PQ-9 in the Q_A_ site binds with high affinity and is located near the monomer-monomer interface. In contrast, the Q_B_ site is occupied by a labile mobile PQ-9 with is peripheral and exchangeable^38^. It is also noteworthy that the non-exchangeable quinones in both PS I_Δ*menB*_ and PS II exhibit bent tails at angles of ∼112° and 91° respectively, unlike the exchangeable ones whose tails are less bent at angles of ∼155° and 133° respectively (**Fig. 7**).

**Fig. 7.**
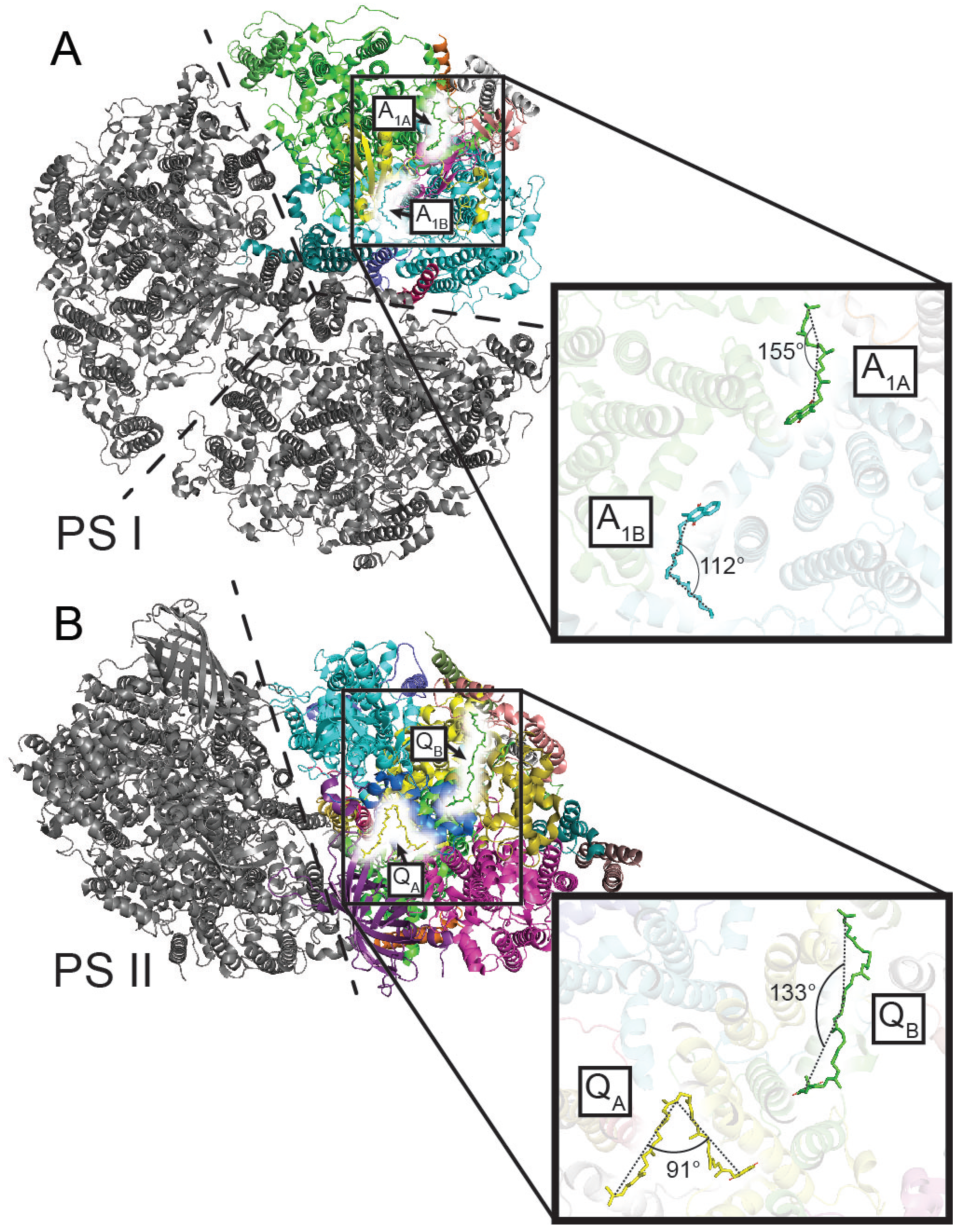
Overlay of the positioning of the quinones within wild-type PS II and PS. **I.** (*A*) Structure of PS I (PDB: 5OY0) with the quinones highlighted in white. (*B*) Structure of PS II (PDB: 3WU2) with the quinones highlighted in white. The insets show approximations of the quinone tail angles.

In summary, we have determined cryo-EM structures of PS I_Δ*menB*_ and PS I_Δ*menB+ENQ*_ isolated from the newly generated *ΔmenB(2024)* strain which delivers structural insight into this important variant. Both the A_1A_ and A_1B_ sites exhibit variable occupancies of different quinone species and/or orientations. By developing and performing a map analysis technique, we deconvoluted the occupancy estimations. Moreover, by employing TR-EPR spectroscopy we confirmed that the binding of the quinones tested here in the *ΔmenB(2024)* strain are identical to the *ΔmenB(2002)* strain. These observations suggest tail lengths of the quinones are an important determinant in their binding affinity, but that protein interactions, both short- and long-range, strongly influence the ability for exchange. This likely applies to all RCs and perhaps all quinone-binding proteins in nature.

## Materials and Methods

### Cell growth and protein purification

The *ΔmenB(2024)* strain was generated and propagated as described in Russell at al.^31^ Cells were grown in medium intensity light at 30 °C in B-HEPES media containing 20 μg mL^−1^ spectinomycin and were harvested by centrifugation, resuspended in 100 mL of buffer (50 mM MES pH 6.5, 15 mM CaCl_2_, 15 mM MgCl_2_), and lysed by 8 passes through a microfluidizer. Cell debris was removed with low-speed centrifugation at 2,000×*g*, and the thylakoid membranes (supernatant) were pelleted by centrifugation at 158,000×*g* for 1 h. Membranes were solubilized at 4 °C in the dark with 1% (w/v) β-dodecylmaltoside (β-DDM) for 1 h. PS I was initially purified on a 5-10% sucrose density gradient 50 mM Tris-HCl buffer (pH 8.3), 0.05% (w/v) β-DDM at 132,000×*g*. The PS I containing band was collected and dialyzed against 200 volumes of 50 mM Tris-HCl buffer (pH 8.3) twice. The sample was then concentrated using an Amicon Stirred Cell concentrator with 100 kDa membrane and ran on a second sucrose gradient (same gradient conditions). The PS I containing band was collected, dialyzed, and concentrated and stored at 4 °C (PS I_Δ*menB*_). To exchange ENQ into the quinone binding sites, 117 µM ENQ was added to PS I_Δ*menB*_ protein stock and incubated overnight at 4 °C. The sample was then washed 3 times with 1000x dilution of ENQ and concentrated to a final concentration of 117 nM ENQ in 50 mM MES pH 6.5, 15 mM CaCl_2_, 15 mM MgCl_2_.

### Negative stain imaging and data processing

4 µL of Δ*menB* PS I sample at 2 µg Chl mL^−1^ was pipetted onto a glow discharged (20 mA for 7 s) Cu Formvar/Carbon 400 mesh electron microscopy grid (Electron Microscopy Sciences) and allowed to incubate at room temperature for 60 s. Liquid was wicked away with a filter paper and the sample side of the grid was touched against a drop of buffer (50 mM MES 6.5 (NaOH), 10 mM MgCl_2_, 10 mM CaCl_2_, 0.02% β-DDM). The liquid was wicked away and touched to a new drop of buffer. This buffer was wicked away, and then this was repeated two times with droplets of 2% uranyl acetate solution. The grid was then floated on a third droplet of 2% uranyl acetate for 60 s before wicking to dry. The negatively stained grids were stored under vacuum for at least 1 h and then imaged on a 120 kV Talos L120C transmission electron microscope. At least 15 micrograph images were collected per sample for data processing. Images were processed using CryoSPARC version 4.7.1^39^. For PS I_Δ*menB*_, 299 particles were selected manually and 2D classification was performed to generate initial classes (*SI Appendix*, **Fig. S4*B***). For PS I_Δ*menB*+ENQ_, 405 particles were selected manually and 2D classification was performed to generate initial classes (*SI Appendix*, **Fig. S4*D***).

### Cryo-EM grid preparation and screening

A glow discharged (25 mA for 30 s) Quantifoil R1.2/1.3 Cu 300-mesh microscopy grid was mounted in a Thermo Fisher Scientific Vitrobot Mark IV system set to 100% humidity and 4 °C. The grid was blotted for 3 seconds with a 0 second wait time and a 0.5 second drain time and plunged into liquid ethane. The sample was then transferred into liquid nitrogen. Cryo-EM screening was performed on an Arctica transmission electron microscope (Thermo Fisher Scientific, FEI). Five micrographs each were collected for PS I_Δ*menB*_ and PS I_Δ*menB*+ENQ_. Example micrograph images are shown in *SI Appendix*, **Fig. S5**. These images were processed using CryoSPARC version 4.7.1^39^. The contrast transfer function was estimated using Ctffind-4^40^. For PS I_Δ*menB*_, 141 particles were selected manually and 2D classification was performed to generate initial classes (*SI Appendix*, **Fig. S5*B***). For PS I_Δ*menB*+ENQ_, 82 particles were selected manually and 2D classification was performed to generate initial classes (*SI Appendix*, **Fig. S5*D***).

### High-resolution cryo-EM data collection and processing

Data was collected on Titan Krios G3 transmission electron microscope (Thermo Fisher Scientific, FEI) operated at 300 kV with a Gatan K3 direct electron detector. The defocus was set to −0.8 to −2.0 μm and the pixel size was 0.417 Å (super-resolution). The imaging filter (GIF) setting was a slit size of 20 eV. The dose was 10.2 e^−^ physical pixel^−1^ s^−1^ with 3.41 s per exposure. Thus, the total dose was 50 e^−^ Å^−2^. Each micrograph movie contained 50 frames. EPU was used to collect 16,258 micrograph movies for PS I_Δ*menB*_ and 16,577 micrographs movies for PS I_Δ*menB*+ENQ_. Data processing was performed using CryoSPARC version 4.7.1^39^. An overview of data processing pipelines can be found in *SI Appendix*, **Fig. S5** and **S6**.

### Model building

The starting model was the previously determined PS I_evolΔ*menB*_ structure (PDB: 9AU4). The model was fit into the map using UCSF Chimera^41^. Manual editing was performed using Coot^42^. Automated refinement and statistical analysis was performed using real_space_refine in the Phenix software suite^43,44^.

### TR-EPR Spectroscopy

TREPR experiments were carried out using Bruker E580 X-band spectrometer equipped with a dielectric resonator and a CF935 cryostat. Time/field datasets were recorded in direct-detection mode without field modulation using diode detection and the signal was digitized using the built in SpecJet digitizer (6 ns per point). The samples were photoexcited at 532 nm using the second harmonic of Continuum Surelite Nd-YAG laser system operating at 10 Hz and ∼4 mJ/pulse. All measurements were carried out at 80 K. Spin-polarized EPR spectra were extracted from the complete time/field datasets in a 500 ns wide time window centered at 250 ns after the laser flash.

## Data, Materials, and Software Availability

The PS I_Δ*menB*_ struture has been deposited in the PDB under (9YL7) (https://www.wwpdb.org/pdb?id=pdb_00009YL7) and EMD under (EMD-73077) (https://www.ebi.ac.uk/emdb/EMD-73077), and the PS I_Δ*menB*+ENQ_ struture has been deposited in the PDB under (9YO1) (https://www.wwpdb.org/pdb?id=pdb_00009YL7) and EMD under (EMD-73249) (https://www.ebi.ac.uk/emdb/EMD-73249) All other data are included in the manuscript and/or SI Appendix.

## ACKNOWLEDGEMENTS

All electron microscopy was performed in the Cryo-EM Research Center (CEMRC) in the Department of Biochemistry at the University of Wisconsin-Madison. We thank the staff of the CEMRC for their helpful guidance. This work was supported by the National Institute of General Medical Sciences of the NIH Award No. R00GM140174 (C.J.G.). The content is solely the responsibility of the authors and does not necessarily represent the official views of the NIH. It was also supported by the Department of Energy, Office of Basic Energy Sciences grants DE-FG02-07ER15903 (K.V.L.), DE-SC0010575 (J.H.G.), and DE-SC0025359 (D.J.V.).

## Supplementary Text

### Text S1. Quantitative cryo-EM map analyses of quinone occupancy

The unmodified electrostatic potential (ESP) maps were converted to charge density (CD) maps according to Poisson equation via negated Laplacian operation using Chimera(1, 2). The experimental CD map features within a box of 20 Å surrounding a quinone were excised and moved to a reference frame in a cubic box with edge length of 20 Å using an initial matrix relating this quinone to the a reference quinone in the center of the reference box, followed by map-tomap alignment to a theoretical CD map at 1.0 Å resolution before being resampled onto the reference CD maps using Chimera. The maps were Fourier inverted to structure factors using CCP4, which were used for quantitative analysis through Fourier synthesis(3, 4). The onedimensional CD map profiles were sampled along 11 consecutive bonds connected to 12 selected atoms of a standard quinone with 201 points for each bond and compared with simulated CD maps with known compositions of quinone mixture (Fig. 5A, S9A).

The CD maps for standard quinones were simulated via ESP in the middle of cubic box with edge length of 20 A at 1.0 A resolution using Peng’s formulations and guesstimate atomic partial charges according to Kollman and colleagues(5–7). The simulated CD maps of two quinone types were then mixed in known proportions with an increment of 25%, i.e., 4:0, 3:2, 2:2, 1:3, and 0:4 ratios, so that the experimental profiles could be compared with the simulated profiles (Fig. 5, S9). An initial B-factor was uniformly set to 5 A2 for all quinone atoms and modified with a ΔB increment of 10 A2 from 0 to 50 A2 so that the effect of local resolution variation on the CD profiles could be considered for composition deconvolution (Fig. 5, S9). Similar simulations were carried out for flipping quinone headgroups.

## Supplementary Figures

**Fig. S1.**
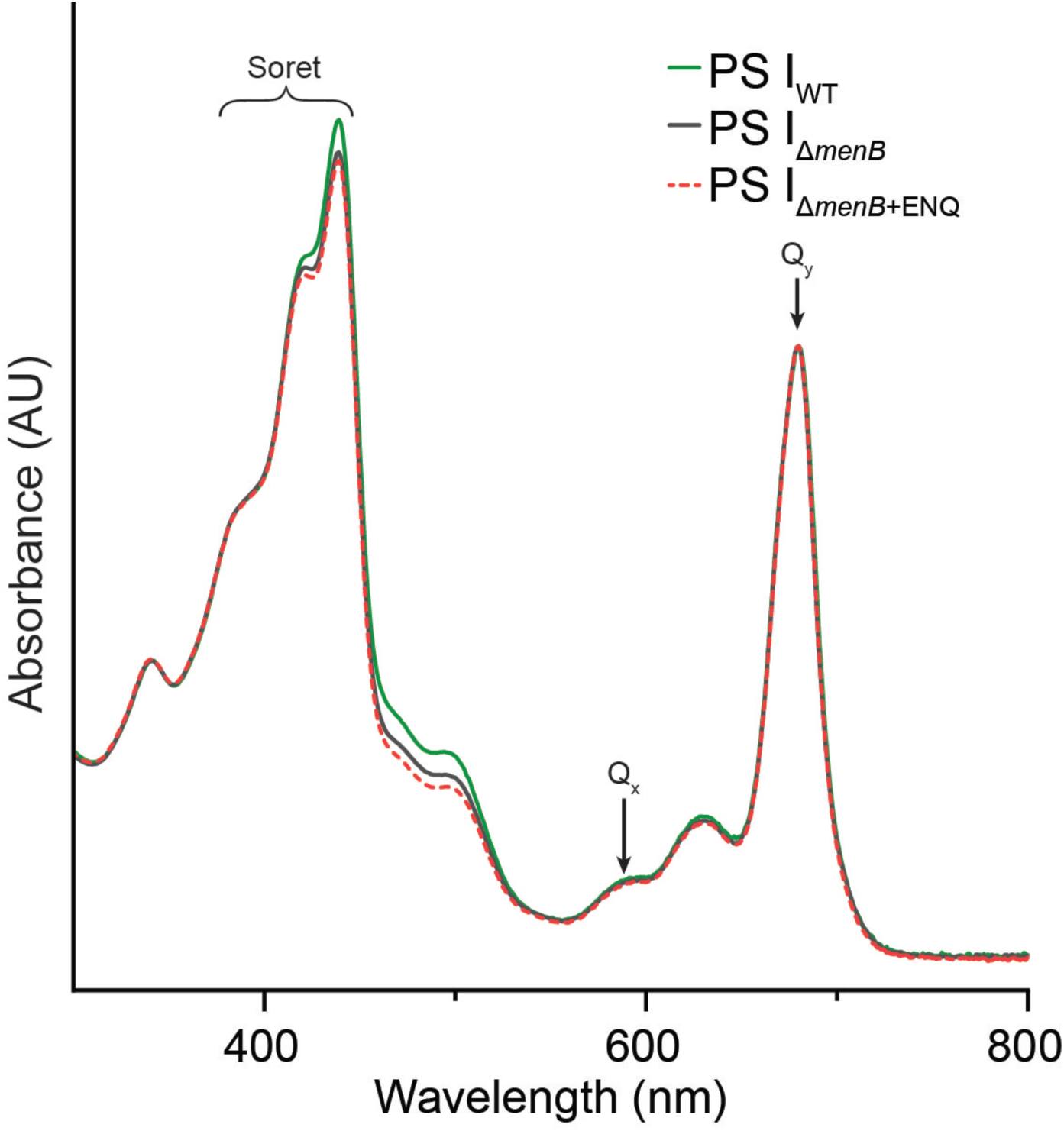
Absorbance spectra of purified PS I samples. Peaks corresponding to Soret transitions, Q_x_, and Q_y_ are labeled. All traces are normalized to the Q_y_ peak.

**Fig. S2.**
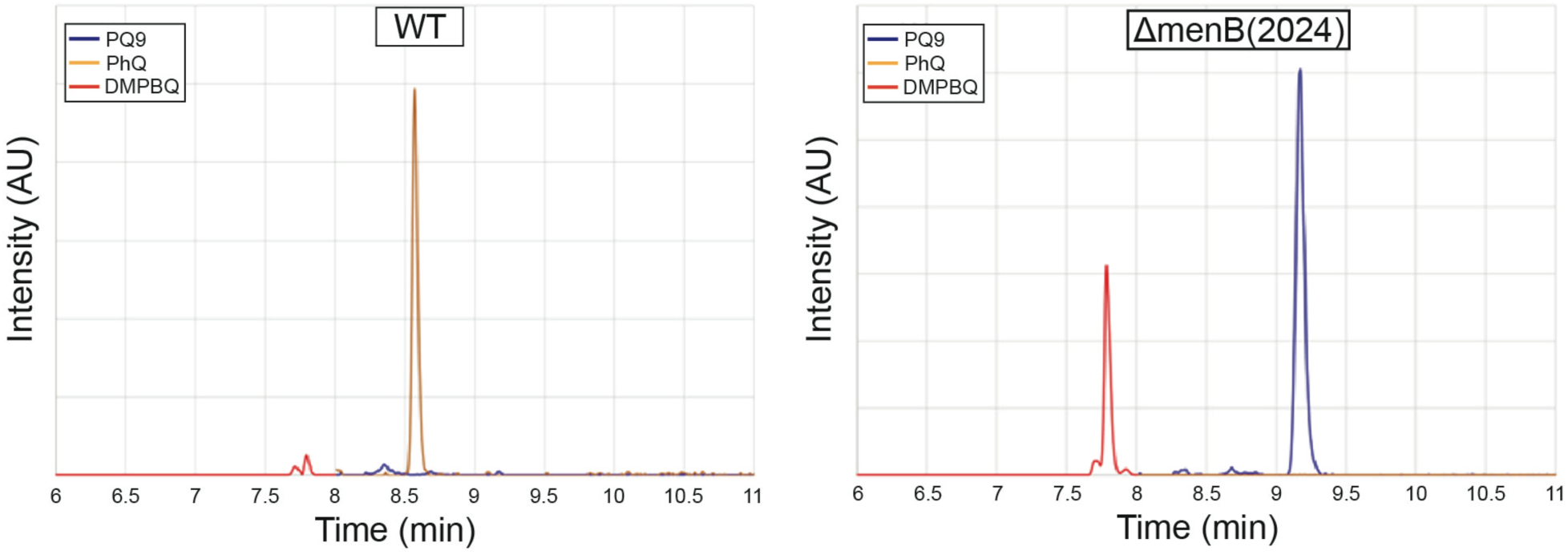
Liquid chromatography/mass spectrometry of quinones. LC/MS ion chromatogram of wild type *Synechocystis* sp. PCC 6803 (WT) and *ΔmenB(2024)* isolated photosystem I. Traces represent species eluting at an *m/z* of 749.6231 (PQ9), 451.3571 (PhQ), and 415.3571 (DMPBQ).

**Fig. S3.**
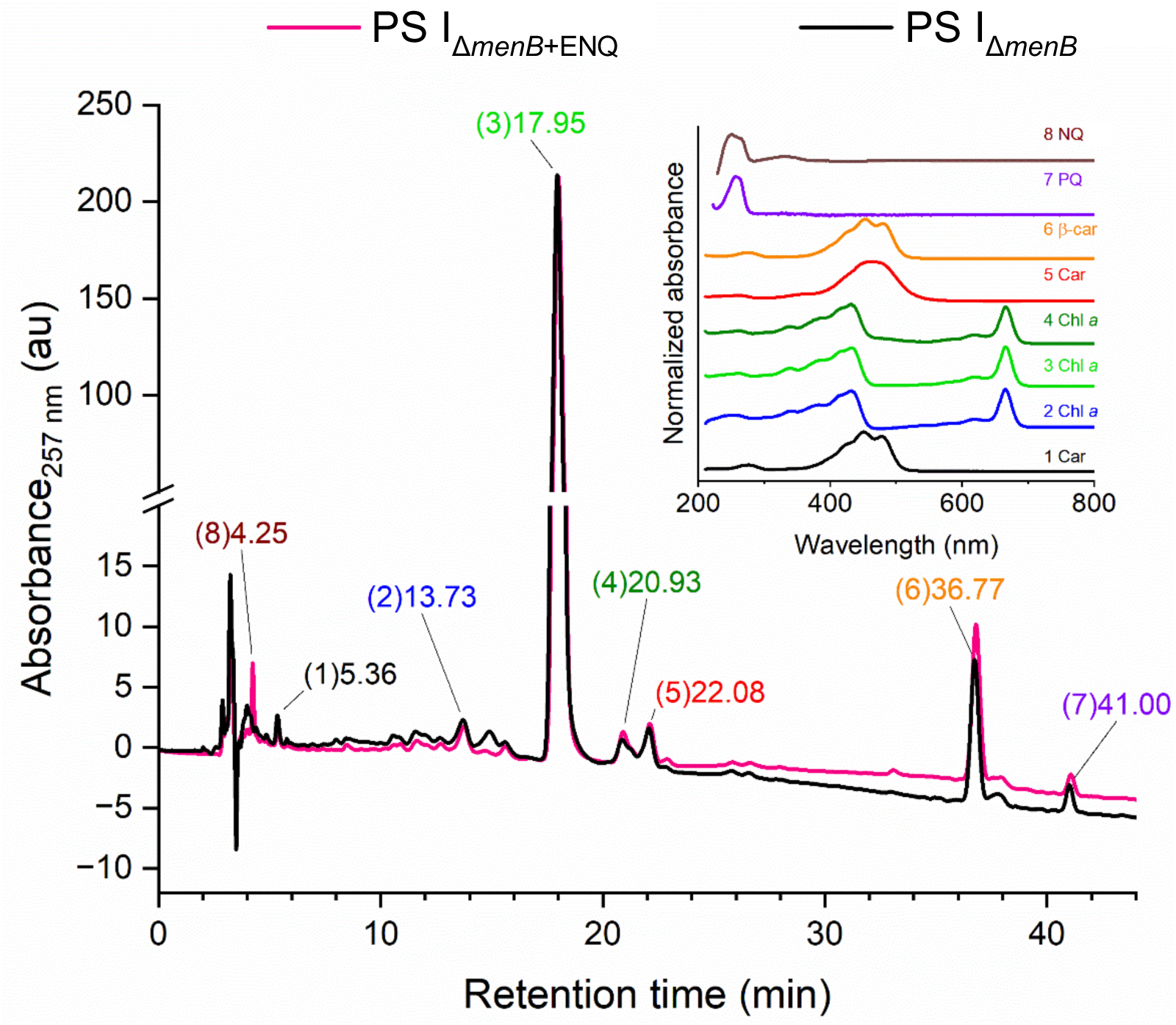
Cofactor analysis of PS I_Δ*menB*_ and PS I_Δ*menB*+ENQ_. HPLC chromatograms were recorded at 257 nm of pigment extracts and normalized to the intensity of the main Chl *a* peak. Peaks are identified according to their absorption spectrum as labeled in the inset.

**Fig. S4.**
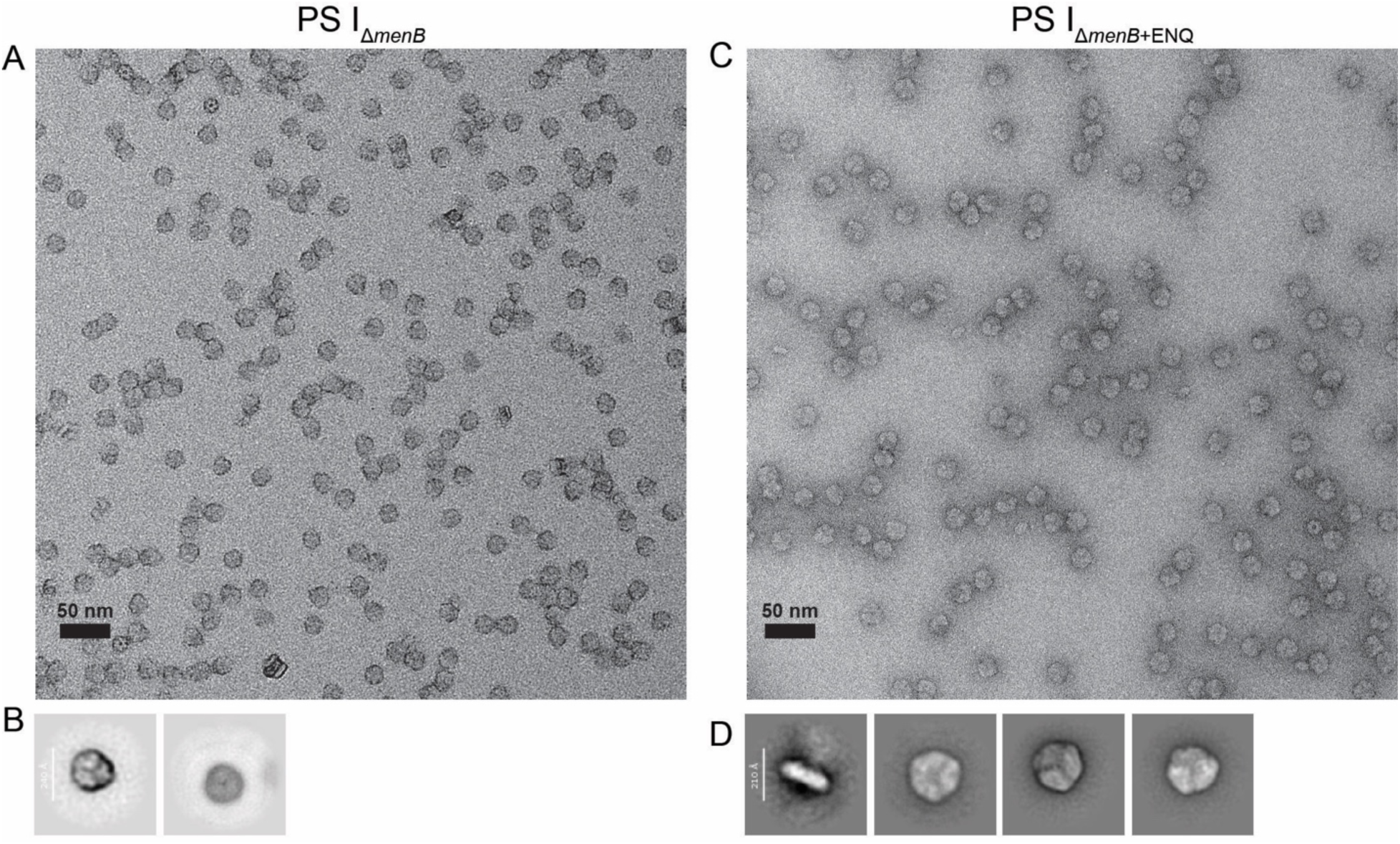
Transmission electron microscopy of negatively stained PS I samples. (*A*) Representative micrograph image of negatively stained PS I_Δ*menB*_ particles. (*B*) 2D classes from 299 particles. (*C*) Representative micrograph image of negatively stained PS I_Δ*menB*+ENQ_ particles. (*D*) 2D classes from 405 particles.

**Fig. S5.**
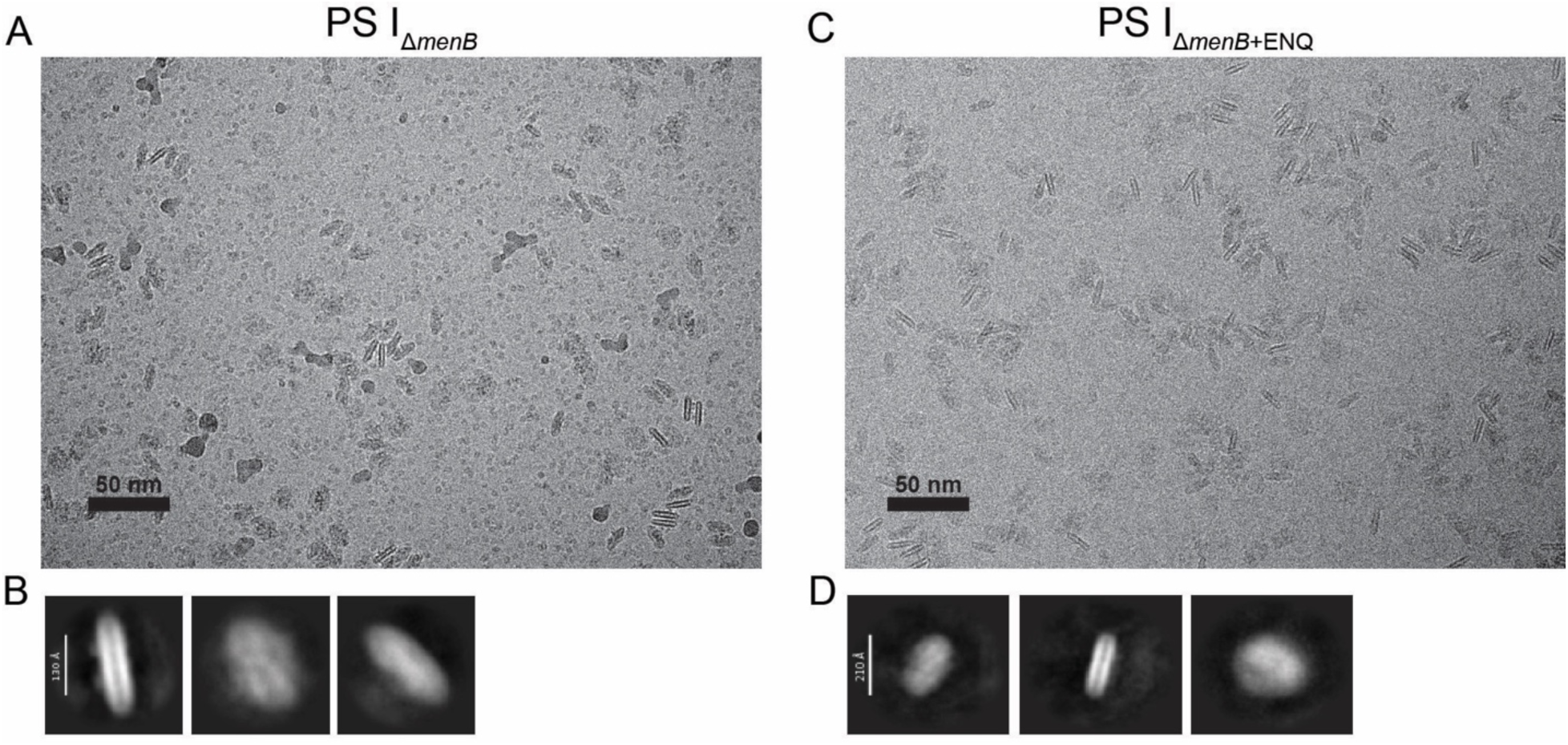
Arctica cryo-EM screening of PS I samples. (*A*) Representative micrograph image of PS I_Δ*menB*_ particles. (*B*) 2D classes from 141 particles. (*C*) Representative micrograph image of PS I_Δ*menB*+ENQ_ particles. (*D*) 2D classes from 82 particles.

**Fig. S6.**
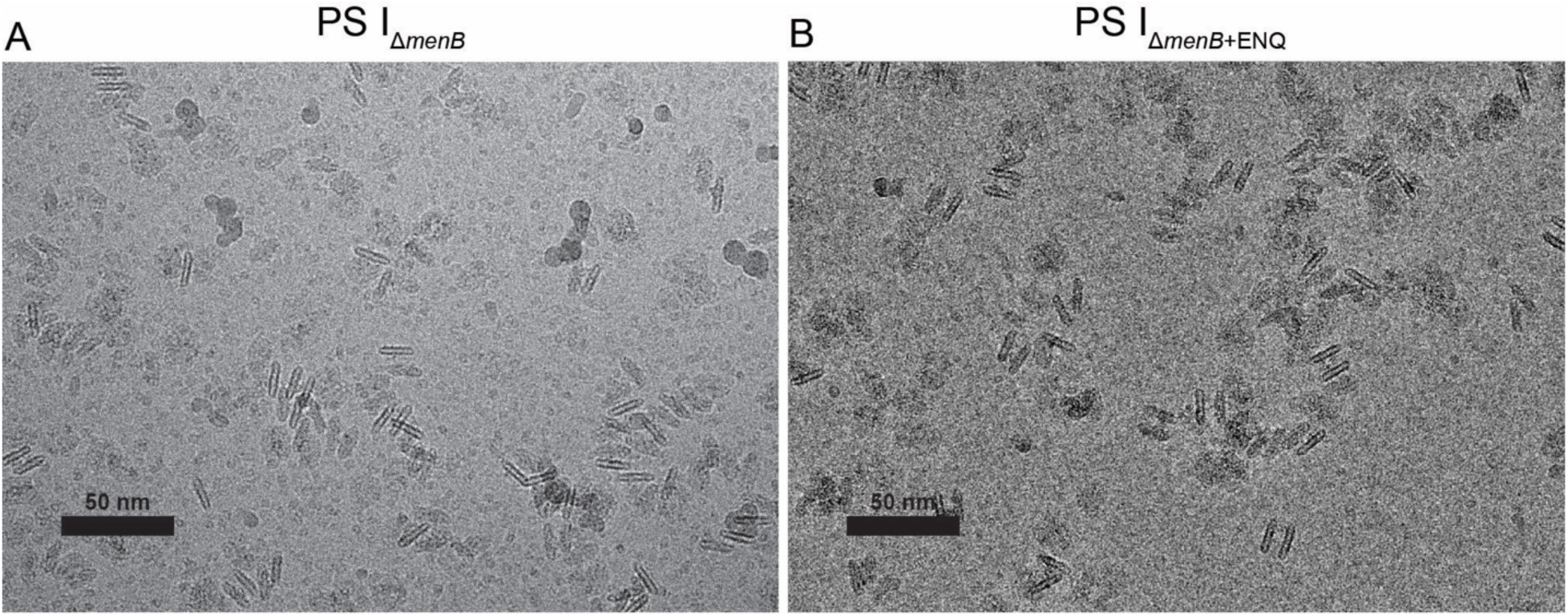
Example micrographs from Krios G3 data collection. (*A*) Representative micrograph image of PS I_Δ*menB*_ particles. (*B*) Representative micrograph image of PS I_Δ*menB*+ENQ_ particles.

**Fig. S7.**
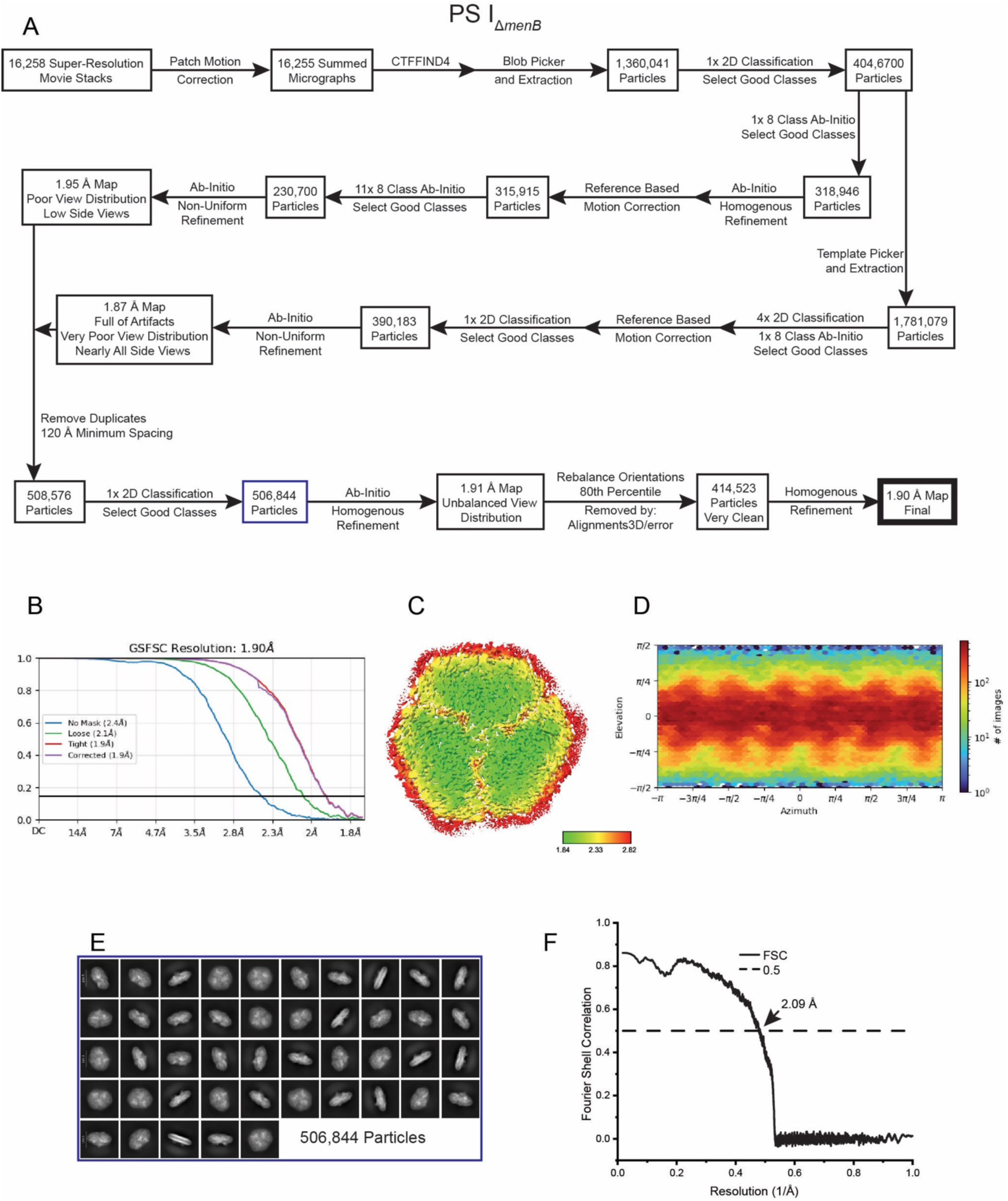
Data processing for the PS I_Δ*menB*_ structure. (*A*) Workflow of cryo-EM data processing in CryoSPARC. (*B*) Gold-Standard Fourier Shell Correlation (GSFSC) of final cryo-EM map. (*C*) Local resolution map, units in Å. (*D*) Final view distribution. (*E*) Last 2D classification in workflow (shown with blue box). (*F*) Map-to-model Fourier Shell Correlation corresponding to sharpened and masked map.

**Fig. S8.**
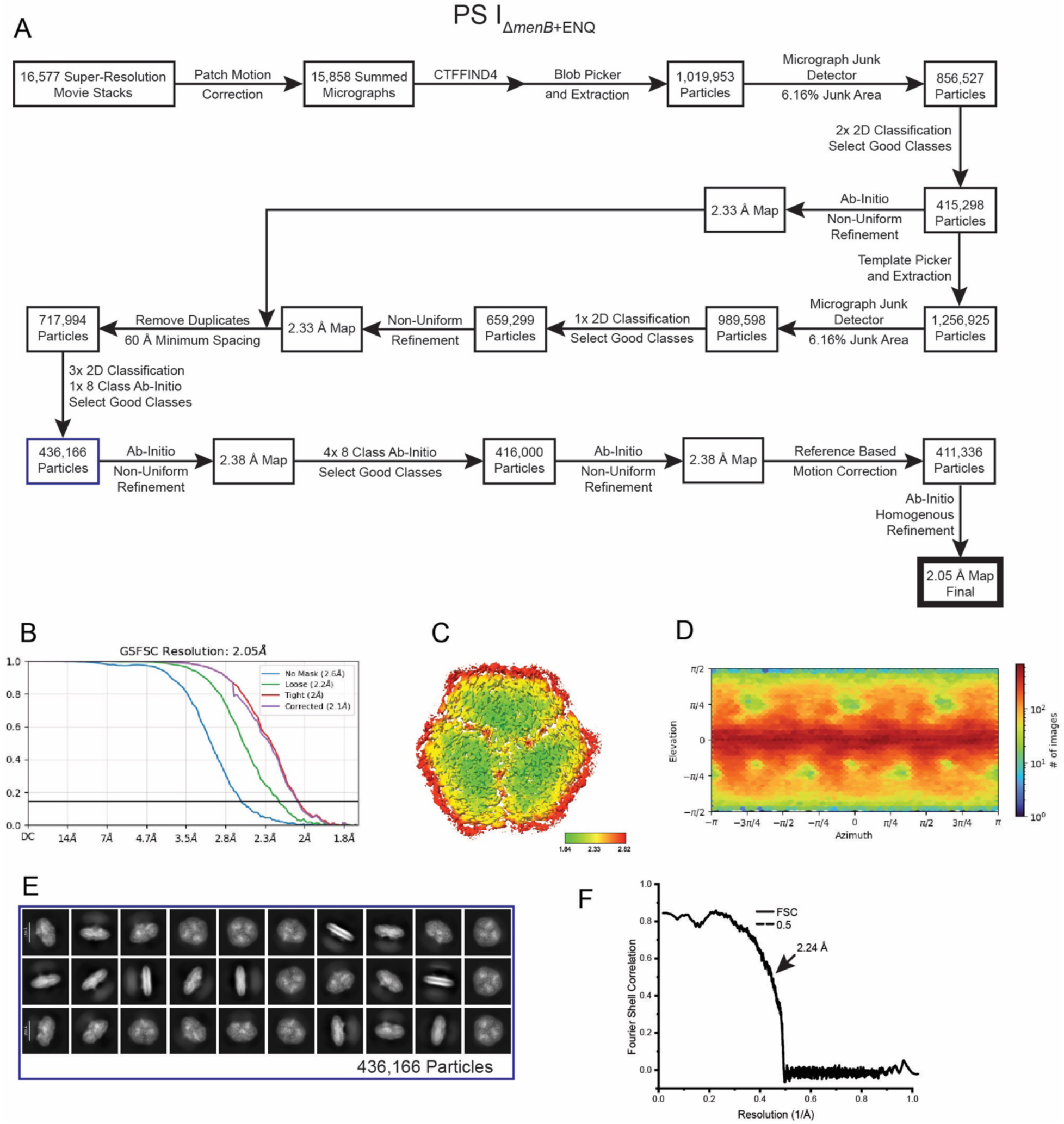
Data processing for the PS I_Δ*menB+EtNQ*_ structure. (*A*) Workflow of cryo-EM data processing in CryoSPARC. (*B*) Gold-Standard Fourier Shell Correlation (GSFSC) of final cryo-EM map. (*C*) Local resolution map, units in Å. (*D*) Final view distribution. (*E*) Last 2D classification in workflow (shown with blue box). (*F*) Map-to-model Fourier Shell Correlation corresponding to sharpened and masked map.

**Fig. S9.**
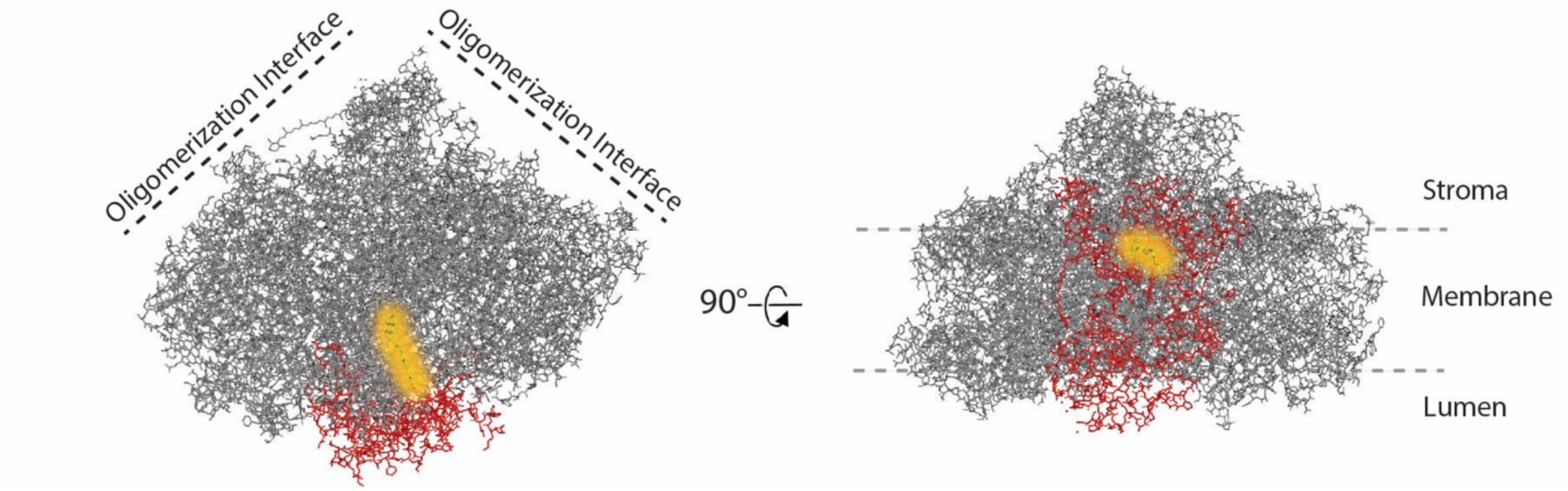
Poorly resolved features in the PS I_Δ*menB*_ structure. The model of PS I reported by Jordan et al. is shown (PDB 1JB0). Features in the PS I_Δ*menB*_ structure that are poorly resolved, likely due to destabilization, are colored in red. The PhQ molecule in the A_1A_ site is colored yellow. This site would be occupied primarily by a PQ-9 in the PS I_Δ*menB*_ structure.

**Fig. S10.**
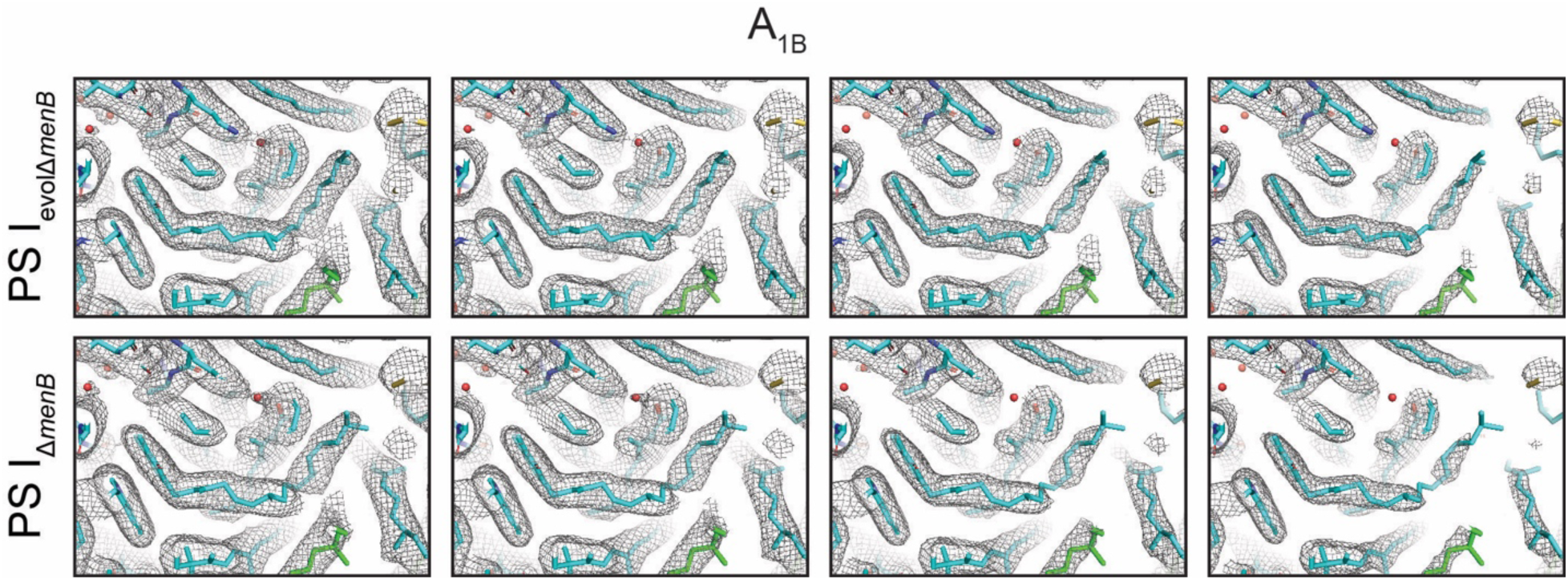
Map of the A_1B_ tails comparing PS I_Δ*menB*_ and PS I_evolΔ*menB*_ shown at different thresholds. The unsharpened map is shown from higher (left) to lower (right) threshold.

**Fig. S11.**
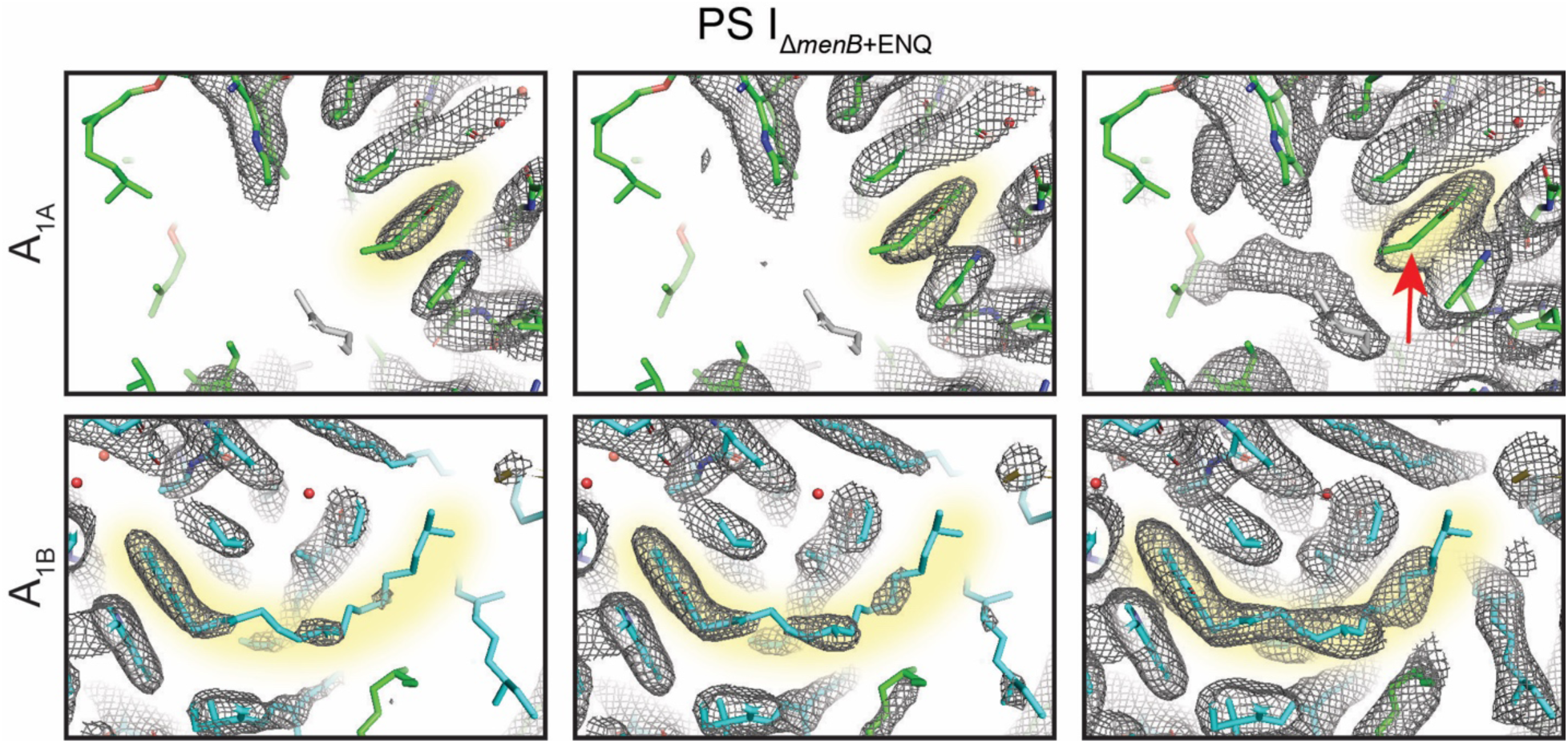
Map of the tails of the quinones in the PS I_Δ*menB+EtNQ*_ structure shown at different thresholds. The unsharpened map is shown from higher (left) to lower (right) threshold. The red arrow indicates the position of the ENQ headgroup.

**Fig. S12.**
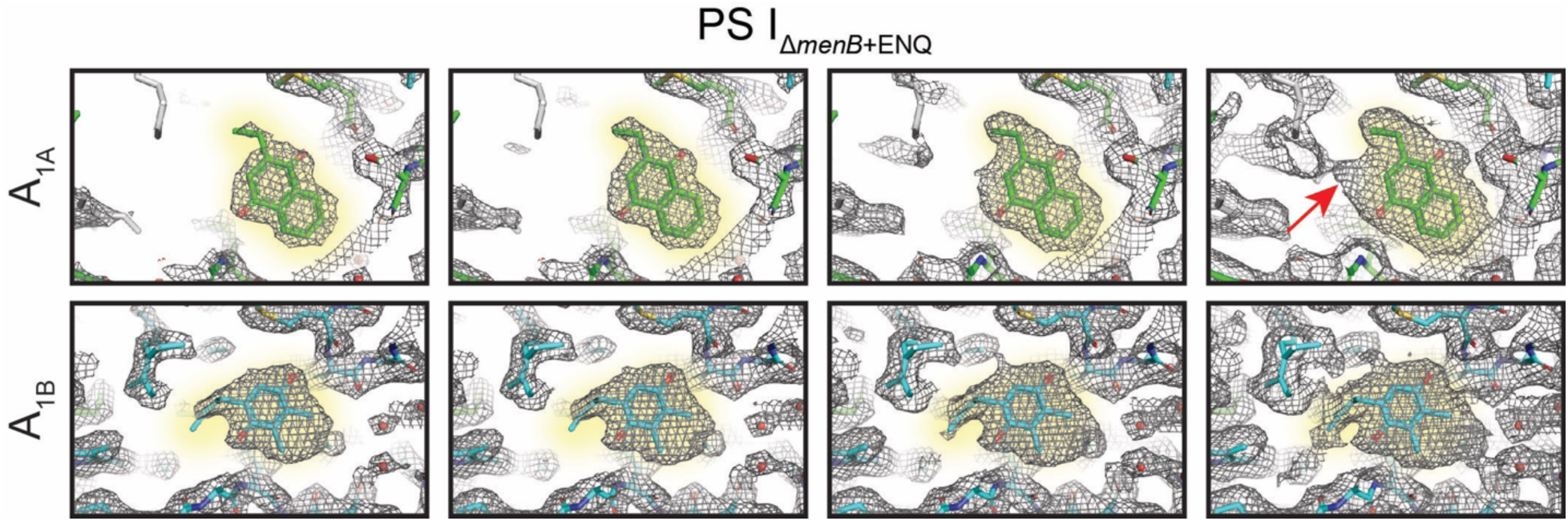
Map of the headgroups of the quinones in the PS I_Δ*menB+EtNQ*_ structure shown at different thresholds. The unsharpened map is shown from higher (left) to lower (right) threshold. The position of the ethyl subunit in the un-flipped ENQ orientation is indicated with a red arrow.

**Fig. S13.**
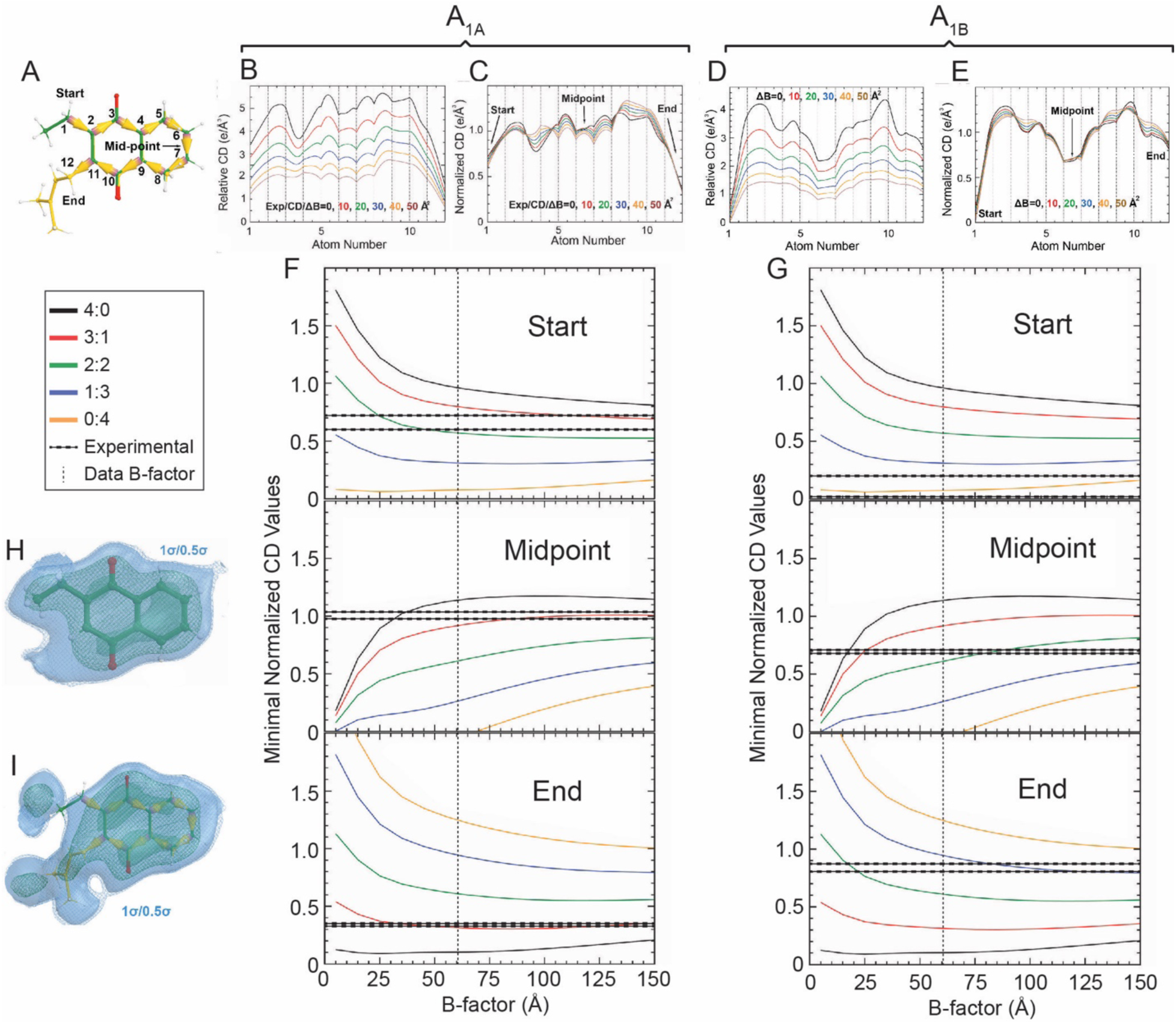
Basis of quantitative analysis of the PS I_Δ*menB+EtNQ*_ cryo-EM map. (*A*) Map of atom numbering to molecular positions, highlighting Start, Midpoint, and End. (*B*) Experimental traces from cryo-EM map at the A_1A_ site converted to charge density and blurred to specific B-factors. (*C*) Normalized experimental traces from cryo-EM map by making the mean value of the entire one-dimensional CD profiles be the unity at the A_1A_ site. (*D*) Experimental traces from cryo-EM map at the A_1B_ site converted to charge density and sharpened to specific B-factors. (*E*) Normalized experimental traces from cryo-EM map at the A_1B_ site. (*F, G*) Modeled traces for the start, midpoint, and end position at varying ratios of quinone identities. Experimental density range shown as two horizontal dashed lines, and experimental B-factor shown and vertical dashed line. Shown for the A_1A_ and A_1B_ sites respectively. (*H, I*) Calculated primary quinone species shown fit into experimental map density at the A_1A_ and A_1B_ sites.

**Fig. S14.**
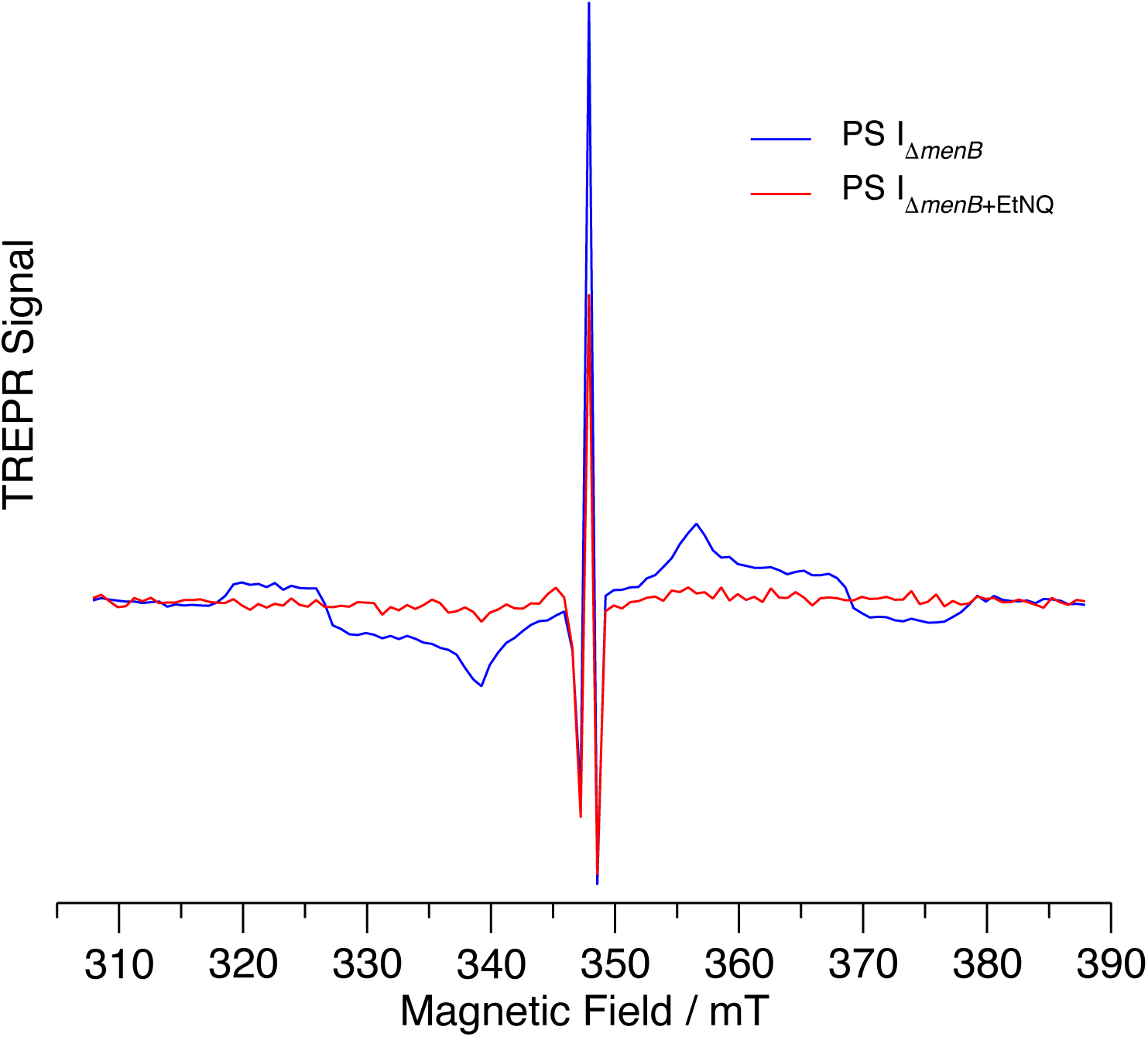
Transient EPR triplet spectrum of PS I_Δ*menB*_ and PS I_Δ*menB*+ENQ_. The broad features in the spectrum of PS I_Δ*menB*_ (blue trace) arise from ^3^P_700_ formed by recombination of P_700_^+^A_0_^−^ when the A_1A_ binding site is empty. These features are absent in the spectrum of PS I_Δ*menB*+ENQ_ (red trace). The sharp peak in the center of the spectra arises from P_700_^+^A_1A_^−^.

**Fig. S15.**
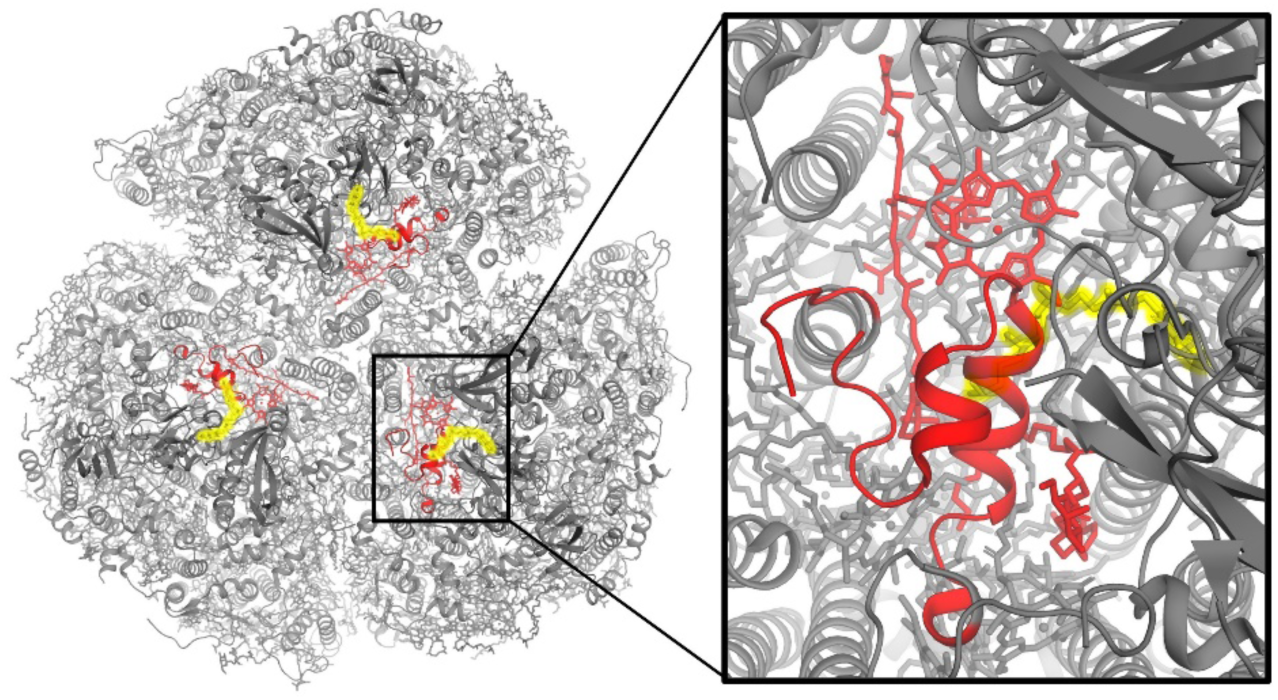
Possible regions of the PS I complex that would need to move if PQ-9 in the A_1B_ site were to exchange. The structure of wild-type PS I is shown with the B-side PhQ highlighted in yellow. Red features of the complex would be expected to move if PQ-9 bound in the A_1B_ site were to exchange.

**Fig. S16.**
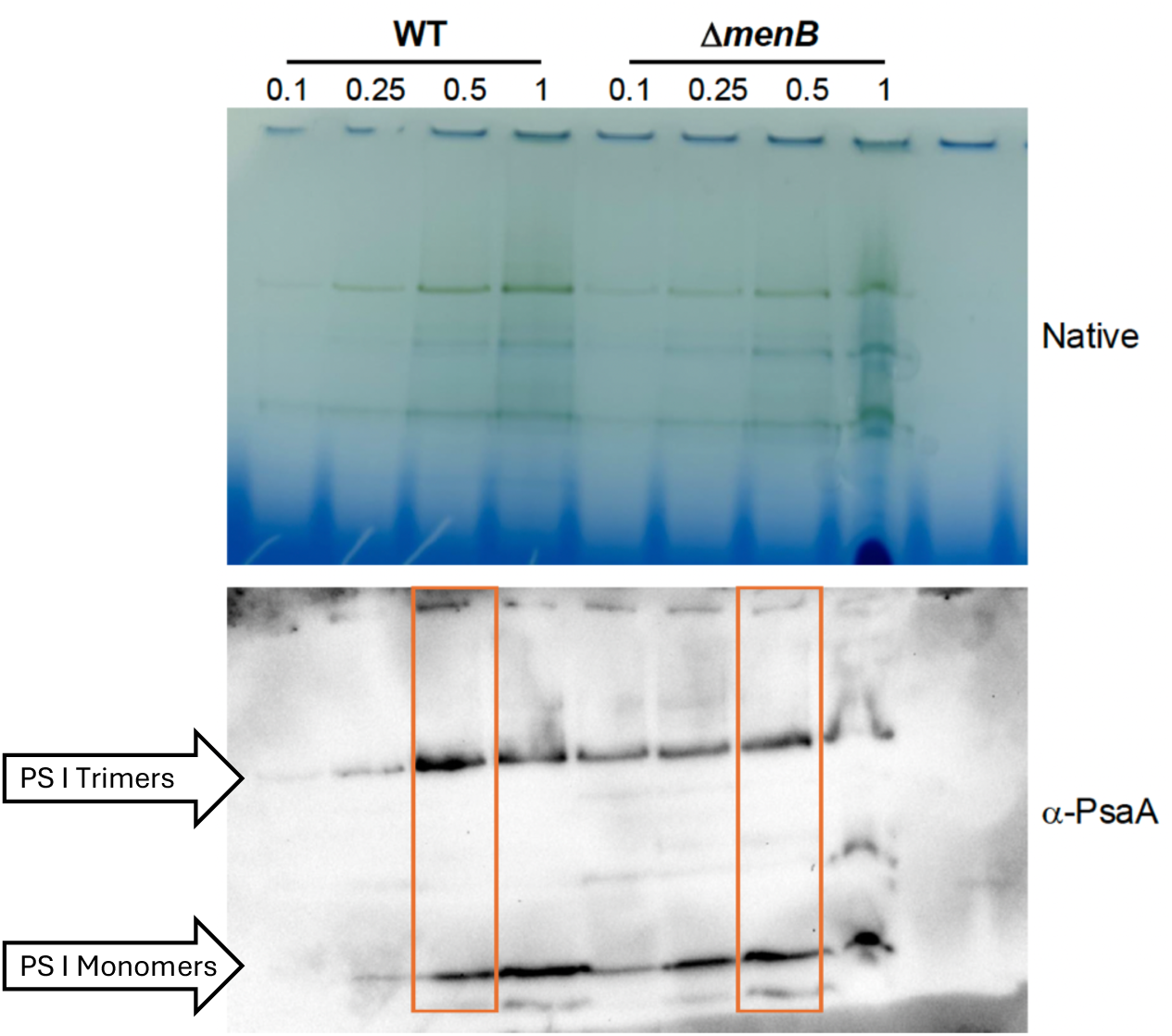
Blue Native PAGE (top) and anti-PsaA Western Blot (bottom) gels. Protein loaded from solubilized thylakoid membranes from *ΔmenB(2024)* and wild type *Synechocystis* sp. PCC 6803 with loading normalized to chlorophyll content. Qualitative measurements drawn from lanes highlighted in orange.

## Supplementary Tables

**Table S1.** Cryo-EM data collection, refinement, and validation statistics for PS I_Δ*menB*_ and PS I_Δ*menB+EtNQ*_.

|  | PS I<sub>ΔmenB</sub> | PS I<sub>ΔmenB+ENQ</sub> |
| --- | --- | --- |
| **Data Collection and Processing** |  |  |
| Magnification | ×105,000 | ×105,000 |
| Voltage (kV) | 300 | 300 |
| Electron Exposure (e<sup>-</sup>/Å<sup>2</sup>) | 50 | 50 |
| Defocus Range (μm) | -0.8 to -2.0 | -0.8 to -2.0 |
| Pixel Size (Å) | 0.834 | 0.834 |
| Symmetry Imposed | C3 | C3 |
| Initial Particle Images (no.) | 1,781,079 | 1,561,354 |
| Final Particle Images (no.) | 414,523 | 411,336 |
| Map Resolution (Å) | 1.90 | 2.05 |
| FSC Threshold | 0.143 | 0.143 |
| **Refinement** |  |  |
| Initial Model Used (PDB) | 9AU4 | 9AU4 |
| Model Resolution (Å) | 2.12 | 2.24 |
| FSC Threshold | 0.5 | 0.5 |
| Map Resolution Range (Å) | 1.87-3.20 | 1.84-4.20 |
| Map-sharpening B Factor (Å<sup>2</sup>) | 49.0 | 61.6 |
| Model Composition |  |  |
| Non-hydrogen Atoms | 73,458 | 71,493 |
| Protein Residues | 6,678 | 6,663 |
| Ligands | 402 | 402 |
| B Factors Mean (Å<sup>2</sup>) |  |  |
| Protein | 23.42 | 39.24 |
| Ligands | 25.32 | 36.63 |
| R.M.S. Deviations |  |  |
| Bond Lengths (Å) | 0.013 | 0.010 |
| Bond Angles (°) | 2.768 | 2.622 |
| **Validation** |  |  |
| MolProbity | 1.78 | 1.79 |
| Clashscore | 7.06 | 5.82 |
| Rotamer Outliers (%) | 0.23 | 0.06 |
| Ramachandran Plot |  |  |
| Favored (%) | 94.28 | 92.51 |
| Allowed (%) | 5.13 | 6.72 |
| Disallowed (%) | 0.59 | 0.77 |

